# Morphological, physiological and metabolic responses of diverse barley inbreds to dry down and moderate drought stress

**DOI:** 10.1101/2024.05.21.595183

**Authors:** Asis Shrestha, Tobias König, Lena Adler Meikle, Philipp Westhoff, Alexander Erban, Benjamin Stich

**Affiliations:** Quantitative Genetics and Genomics of Plants, Heinrich-Heine-University, Düsseldorf; Plant Biochemistry, Heinrich-Heine-University, Düsseldorf; The Cluster of Excellence on Plant Sciences, Heinrich-Heine-University, Düsseldorf; Max Planck Institute of Molecular Plant Physiology, Potsdam; Institute for Breeding Research on Agricultural Crops, Julius Kühn-Institute, Federal Research Center for Cultivated Plants, Sanitz

**Keywords:** drought survival, drought stress tolerance, photosystem II efficiency, proton motive force, metabolite accumulation, stomata density

## Abstract

Drought stress alters the metabolic activity, physiological processes, and plant growth and such responses might differ with the intensity of stress. We evaluated the genotypic diversity on plant morphology, photosynthetic responses, metabolite shift and their relationship in diverse barley inbreds under dry down (DD) and moderate drought (MD) stress using 23 genetically diverse parental inbreds of genetic mapping population of barley. MD stress caused a strong growth reduction, while DD stress triggered inhibition of photosynthetic health. We observed that the induced changes occurred in a genotype-dependent manner. Compared to control conditions, the metabolism of simple sugars and polyhydroxy acids increased in MD and DD, while the maximum accumulation of amino acid, lipids and phosphates occurred in DD stress. Accumulation of hexose and metabolites with unknown classification was the metabolic signature of drought tolerant inbreds. The inbreds tolerant to MD originated from the temperate regions while those tolerant to both MD and DD came from semi-arid regions. Low stomata density, reduced water loss and retarded growth under drought stress were the key features of inbreds with better survival capacity under severe dehydration. We identified drought tolerant barley inbreds and our study offers resources for future genetic research on various drought tolerance strategies.

## Introduction

Agricultural drought is the period during a cropping season when the rainfall and soil moisture cannot meet the evapotranspiration demand of the crop to maintain optimum growth and production (Trenberth *et al*., 2014). Short-term and prolonged dry periods occurred in several historical instances, which were primarily linked to tropical oceanic temperature fluctuations (Wang *et al*., 2018*a*). However, the global change in temperature because of greenhouse gas emissions and land-use change are the key drivers of the increase in frequency and intensity of drought events in the last two decades, causing severe crop damage and economic loss (Hari *et al*., 2020).

Acute water stress does not occur suddenly in nature (Verslues *et al*., 2023). Typically, drought stress is mild during the early stages of water stress. In later stages, plants experience dehydration stress when the water levels in the topsoil are strongly reduced (Verslues *et al*., 2023) which causes severe wilting (Izanloo *et al*., 2008). In addition, the intensity of drought stress might vary with geographical location. For instance, the extent of water limitation in the temperate regions are typically mild. In these regions, the moderate stress rarely results in plant death, instead it limits growth and biomass production (Skirycz *et al*., 2011). However, in the scientific literature, drought stress is assessed predominantly under severe drought scenarios, in the following termed dry down (DD) stress, by terminating water supply and assessing survival (Skirycz *et al*., 2011; Verslues *et al*., 2023). High survival under dry down stress is linked to water conservation strategies, which might lead to growth penalty under moderate drought (MD) stress. However, only little information is available regarding the genetic diversity of tradeoffs between survival mechanism and enhanced plant growth under MD stress (Kooyers, 2015). Furthermore, comprehensive studies on drought-induced change in plant metabolism, morphology and physiology using an extensive set of diverse genetic materials in crops are very rare (Skirycz *et al*., 2011).

Plants undergo morphological and physiological adjustments during drought stress (Yoshida *et al*., 2015) and such adjustments vary with drought intensity (Kooyers, 2015). If the transpiration rate exceeds the water uptake, the initial response of most plants is to avoid water-loss by closing their stomata (Mata and Lamattina, 2001). Additionally, drought stress induces the production of reactive oxygen species, which damages macromolecules and the photosynthetic apparatus (Vendruscolo *et al*., 2007; Shrestha *et al*., 2022). The majority of the available drought studies involving multiple genotypes scores change in plant morphology and leaf greenness, whereas genetic variation of photosynthetic parameters is typically not assessed (Teulat *et al*., 1998, 2001; Baum *et al*., 2003; Talamè *et al*., 2007; Arifuzzaman *et al*., 2014; Sallam *et al*., 2015).

Besides changes in plant growth and photosynthetic parameters, drought stress alters plant metabolism (Templer *et al*., 2017). Accumulation of soluble sugars and amino acids are common response of plant species and such changes might have osmoprotective role under drought stress (Lawas *et al*., 2019). Likewise, drought-induced change in polyols and low molecular organic acids such as ascorbate with antioxidant properties were also reported for barley, wheat and rice (Hochberg *et al*., 2013). The components of membrane lipids and epicuticular wax are other primary metabolites known to accumulate under drought stress (Bondada *et al*., 1996). Previous studies describing changes in metabolite profiles under terminal drought or moderate stress comprised a limited number of genotypes (Bowne *et al*., 2012*a*; Hochberg *et al*., 2013; Todaka *et al*., 2017*b*; Lawas *et al*., 2019; AbdElgawad *et al*., 2020*a*). However, there are no previous reports on change in plant metabolism under different levels of drought stress in barley using a large number of diverse genotypes.

In the current study, we used barley as model to understand the above-mentioned aspects. Barley is one of the major crops grown across the globe from the highlands of Nepal and Ethiopia, in the Tibetan plateau to warm and dry areas in Israel, Jordan and Morocco (Badr *et al*., 2000). Therefore, it offers a unique resource for studying diverse abiotic stresses, including drought stress (Dawson *et al*., 2015).

The objectives of our study were to,

1. evaluate the change in plant morphology, the physiological response, especially photosynthetic parameters and metabolite profiling of diverse barley inbreds under DD as well as MD stress,
2. identify the genotypic diversity of response to DD and MD stress, and
3. understand the general relationship among metabolite shift, morphological and physiological changes under DD and MD stress.

## Materials and Methods

### Plant material, growth conditions and stress treatment

We performed drought experiments in the greenhouse using the 23 parental inbred lines (Table S1) of the double round robin population (Casale *et al*., 2022). Seeds of the 23 barley inbreds were sown in 96 well trays filled with peat-based potting mixture. The trays were placed in a cold room at 4 °C for 48 h. Then, the trays were transferred to the greenhouse. Uniform seedlings were transferred to 1.5 l pots 2 d after germination. Each parental inbred received three treatment levels control, moderated drought (MD), and dry down (DD) stress. We used three replicates per inbred per treatment in two independent experiments. Each treatment had five trays, each accommodating 15 pots. The chance of an inbred being assigned on a tray was completely randomized. The trays were shuffled across the greenhouse every second day to avoid bias due to light distribution and positioning. During the establishment phase (until two weeks after transplanting), the pots were watered to maintain 30% volumetric moisture content (VMC). After two weeks, pots of the control treatment received regular watering keeping 30% VMC. For the MD treatment, water stress was applied by withholding watering until the VMC reached 15%. After that, the VMC in the pots exposed to MD were daily watered to maintain 15% VMC. The DD treatment involved monitoring the moisture content in each pot to control the rate of dehydration. The pots with VMC below the mean of those exposed to DD were watered to balance the variation in moisture content under dehydration treatment. It took 6 d after dehydration treatment for the moisture content to stabilize across the pots (7.2 ± 2.4% in experiment 1 and 9.2 ± 1.8% in experiment 2). Plants were exposed to DD stress for 12 d and then were re-watered to 30% VMC. The moisture reading and weather data during the experiments are provided in Figure S1.

### Morphological and physiological characterization

Non-destructive measurements of photosynthesis-related parameters were performed using a hand-held multi-sensor, MultispeQ V2.0 (PhotosynQ Inc, California, USA) (Kuhlgert *et al*., 2016). Fluorescence yield and absorbance change were recorded at wavelengths ranging from 450 nm to 950 nm using the manufacturer-designed protocol (Photosynthesis RIDES 2.0). The protocol allows to determine several photosynthetic parameters, including photochemical yield in PS II (Φ_II_), non-photochemical quenching (NPQ) and proton flux across thylakoid membrane. In addition, the protocol also estimates relative chlorophyll (RC) content and leaf thickness (LT). The data were recorded from the middle (lengthwise) of the first fully expanded leaf from the top. Similarly, morphological parameters such as plant height (PH) and tiller number (TN) were evaluated during the stress period. Photosynthesis and morphological data were collected at 1 d, 4 d and 7 d after exposition to DD treatment and 1 d, 4 d, 7 d, 11 d, 14 d, 18 d, 21 d, 25 d and 28 d after exposition to MD stress.

### Relative water content

Leaf water status was estimated as relative water content (RWC) according to Shrestha *et al*. (2022). For RWC measurement, two pieces of 1 cm leaf sections were excised from the middle of the second fully expanded leaf of the tallest tiller, and the fresh weight was recorded (FW). Then, the leaf sections were placed in a centrifuge tube (Falcon® 15 mL) filled with 10ml deionized water for 24 h at room temperature. The leaf sections were removed from the falcon tube, and excess water was wiped with a paper towel before taking the turgor weight (TW). Dry weight was recorded after oven drying at 70^°^C for 24 h. RWC was estimated as (FW-DW)/(TW-DW)*100.

### Metabolite analysis

We collected the first fully expanded leaf from the top from the two tallest tillers from each plant. The samples were snap frozen in liquid nitrogen. Frozen samples were stored at −80 °C before further processing. Leaf sampling was done 7 d and 12 d after the start of DD (VMC of 5.3 ± 1.8% in experiment 1 and 6.8 ± 1.1% in experiment 2) and MD stress (VMC of 12.5 ± 2.4% in repetition 1 and 13.7 ± 2.7% in repetition 2), respectively. For technical reasons, we pooled the samples from the plants under controlled conditions, 7 d (leaf sampling according to DD stress) and 12 d (leaf sampling according to MD stress) after the start of stress treatment. Therefore, we have a single control sample for the metabolite pool under well-watered conditions resulting in 414 samples across the three treatments. Metabolite profiling of each sample was done following gas chromatograph-mass spectrometry (Lisec *et al*., 2006) as described in Wu *et al*. (2022). The mass-spectra were annotated using the Golm Metabolome Database mass-spectral library (Kopka *et al*., 2005) and relative quantification as mentioned in Wu *et al*. (2022).

### Data processing and statistical analysis

Because the time of day might affect the photosynthesis-related processes, the start and end of the measurements were divided into constant time windows of 30 minutes. For instance, the data collected between 8:00 and 8:30 were grouped into time window one and every progressive half-hour was grouped to a subsequent time interval. Hence, for all the photosynthetic parameters, the effect due to experimental repetition, time window and tray could be adjusted. Furthermore, the light intensity measured as photosynthetic active radiation (PAR) showed a significant linear correlation with the derived photosynthetic efficiency parameters (Figure S2). Therefore, the effect of light intensity was adjusted using PAR as an additive covariate in the linear model for estimating the adjusted entry means of the photosynthetic parameters (Gao *et al*., 2024). Statistical significance test, heritability estimate and adjusted entry mean (AEM) were computed from data collected from control and MD stress 1 d to 28 d together. Same was true for data collected from 1 d to 7 d from control and DD stress. For photosynthetic parameters, the analysis of variance (ANOVA) was performed using the following model twice, one-time control and MD, as well as control and DD together:

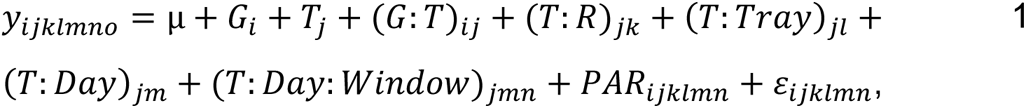

where *y*_*ijklmno*_was the phenotypic observation, (*G*_*i*_) was the effect of genotype, (*T*_*j*_) was the effect of treatment, (*G*: *T*)_*ij*_ was the effect of genotype by treatment interaction, (*T*: *R*)_*jk*_ was the effect of experiment nested within treatment, (*T*: *Tray*)_*jm*_ was the effect of tray nested within treatment, (*T*: *Day*)_*jn*_ effect of day nested within treatment, and (*T*: *Day*: *Window*)_*jno*_ was the effect of time window nested within day after the stress treatment nested within treatment, and *PAR*_*ijklmn*_ was the effect of PAR on each measurement.

Next, we estimated the adjusted entry means (AEM) for each genotype for each treatment condition based on the following general linear model:

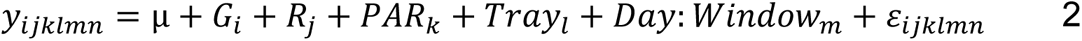

We calculated the AEM for morphological parameters using a modified version of model 2, where we removed the effect of time window and PAR from the model. Because we performed metabolite profiling in samples collected at a single time point and a single control reference sample,

i. the statistical analysis for metabolite data for control, DD and MD were performed together using model 1 and,
ii. the AEM of metabolites of inbreds at different treatment conditions was calculated using model 2,

where the effect of day, time window and PAR were removed from the models. Next, we estimated the repeatability of the parameters per treatment where genotype was used as a random variable in model 2. Repeatability was calculated for each treatment condition using the following formula:

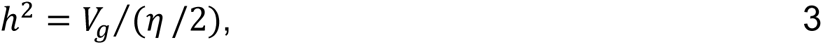

where *V_g_* was the genotypic variance and *η* the average standard error of all pairs of contrasts among genotypes (Piepho and Möhring, 2007). Finally, we interpreted the data in terms of stress response indexes (SRI) calculated as ratios of trait values under stress conditions divided by control conditions. The error associated with the SRI was calculated based on the rules of error propagation.

### Evaluation of stomata density

The 23 parental inbreds were grown in an independent growth chamber experiment for two weeks. Then, we prepared epidermal imprints using nail varnish from the second fully expanded leaf from the bottom. Imprints were prepared from the adaxial and abaxial surface of the leaf at three different positions in the leaf blade: top, middle and the bottom third amounting to six imprints per leaf. The images of epidermal imprints were obtained using a stereo microscope Nikon SMZ18 equipped with a Nikon DS-Fi2 camera at 6X magnification. The size of the image was 1020*1024 pixels. The number of stomata was manually counted using cell-counter plugin in ImageJ. Stomata density (SD) was expressed as the number of stomata mm^-2^. The experiment was performed twice with six biological replicates each following a completely randomized design. The ANOVA for SD was performed using the following equation:

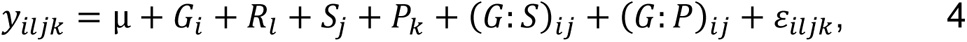

where *y*_*iljk*_ was the observation of SD, *G*_*i*_ was the effect of genotype, *R*_*l*_ was the effect of experiment, *S*_*j*_ was the effect of leaf surface (adaxial or abaxial), *P*_*k*_ was the effect of position on the leaf, (*G*: *S*)_*ij*_ was the effect of genotype by surface interaction, (*G*: *P*)_*ij*_ was the effect of genotype by position. Because we observed a significant (p < 0.001) genotype by position interaction (Table S2), we estimated AEM and *h*^2^ of SD for each position on the leaf separately using model 4 (with the effect of leaf surface removed from the model) and formula 3, respectively.

## Results

### Effect of drought stress on morphological and photosynthetic parameters

The RWC of the leaves of the barley genotypes was assessed under DD and MD conditions at 7 d and 12 d after the start of stress. We observed significant (p < 0.05) treatment and genotype by treatment interaction effects for RWC when analyzing control and MD together as well as control and DD stress together (Table 1). Averaged across 23 parental inbreds, RWC was around one third lower in DD (42%) than MD (71%) stress, illustrating the milder water stress under MD stress conditions compared to DD (Figure S3). Next, we evaluated several growth-related and photosynthetic parameters in these inbreds under well-watered, DD and MD stress. Data were collected until 7 d and 28 d after the start of stress under DD and MD stress, respectively. We observed a significant (p < 0.001) genotype as well as treatment effect on PH, TN and LT under DD and MD stress when analyzing control and MD together as well as control and DD stress together. A significant (p < 0.05) genotype by treatment interaction was also detected for all growth-related parameters under MD and DD (except TN) stress when analyzing control and MD together as well as control and DD stress together (Table 1). The decrease in PH and TN compared to the control condition was more pronounced in MD than in DD stress (Figures S4-S5). Leaf thickness also decreased in DD and MD compared to control conditions, indicating lower moisture content under water stress (Figure S6). The observations across the inbreds were highly repeatable for PH (*h^2^* from 0.91 to 0.99), TN (*h^2^* from 0.79 to 0.95) and LT (*h^2^* from 0.85 to 0.96) under control, DD and MD stress (Table S3).

**Table 1:**
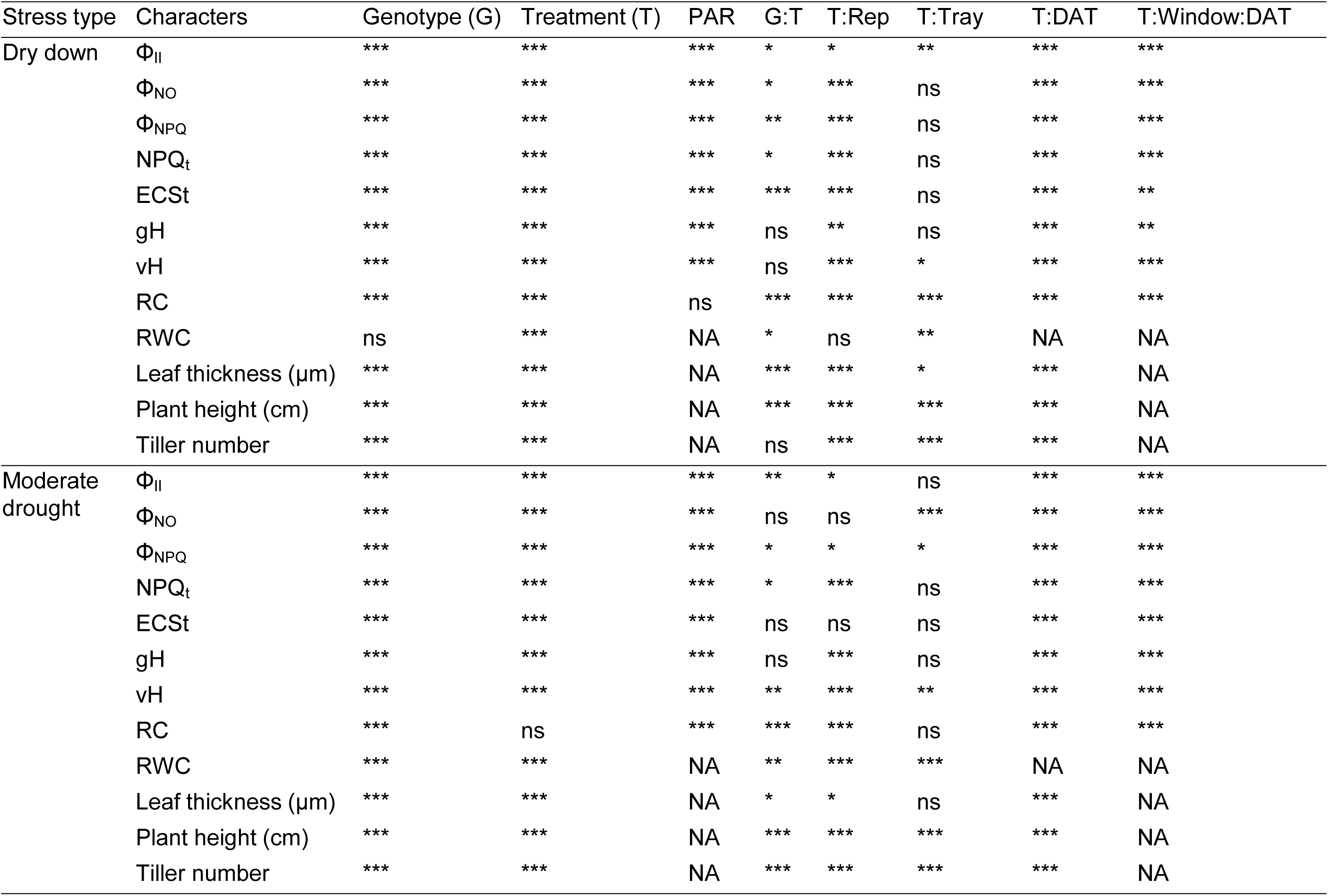
Significance of experimental factors and their interactions on the variation of morphological, physiological and photosynthetic parameters. The experimental factors include genotype (T), treatment (T), two independent experiments (Rep), trays to organize the pots (Tray), days after the start of drought stress treatment (DAT) and time window within the day of measurement (Window). Photosynthetic active radiation (PAR) was fitted as additive co-variable in the linear model. Asterisks indicate the level of significance (***, p ≤ 0.001; **, p ≤ 0.01; *, p ≤ 0.05; ns, p > 0.05; NA, not applicable). Abbreviations: non-photochemical quenching of chlorophyll fluorescence in photosystem II (Φ_NPQ_), NPQ corrected for dark adapted reaction (NPQ_t_), quantum yield of photosystem II (Φ_II_), quantum yield of non-regulatory energy dissipation (Φ_NO_), relative chlorophyll index (RC), magnitude of electrochromic shift (ECS_t_), proton conductivity (gH^+^) and steady state proton flux (vH^+^).

The genotypic variance and drought stress treatment effect were also statistically significant (p < 0.001) for all photosynthetic parameters (Ecs_t_, gH^+^, vH^+^, Φ_II_, Φ_NO_, Φ_NPQ_ and NPQ_t_) and RC measured using MultispeQ when analyzing control and MD together as well as control and DD stress together (Table 1). A significant (p < 0.05 to 0.001) genotype by treatment interaction was observed for vH^+^, Φ_II_, Φ_NO_, Φ_NPQ_, NPQ_t_ and RC for control and MD analyzed together and for Ecs_t_, Φ_II_, Φ_NO_, Φ_NPQ_, NPQ_t_ and RC control and DD analyzed together (Table 1). The *h^2^* was highest for RC compared to gH^+^, vH^+^, Φ_II_, Φ_NO_, Φ_NPQ_ and NPQ_t_ across all growing conditions. There was a minimal difference in the *h^2^* of photosynthetic parameters between control and MD, while the difference in *h^2^* between control and DD was very high, especially for Φ_II_, Φ_NO_, Φ_NPQ_ and NPQ_t_ (Table S3). One reason for this finding is that the number of observations from a plant was threefold higher in the control dataset used in MD (nine observations) than DD (three observations) because of the shorter duration of the experiment in the former than the latter. In conclusion, the phenotypic values of gH^+^, Φ_II_ and Φ_NO_ decreased under DD and MD compared to control condition while vH^+^, Ecs_t_, Φ_NPQ_ and NPQ_t_ increased under both stresses (Figure S8-S14).

### Genotype specific response to dry down and moderate drought stress

First, we performed principal component (PC) analysis (PCA) based on the adjusted entry means of the three treatments separately. Based on the phenotypic diversity, the inbreds originating from Europe clustered together, while those from South Asia, West Asia and Northern Africa grouped together under control conditions (Figure S15A). Likewise, the inbreds from East Asia and Lakhan (from South Asia) and those from the Americas formed separate clusters (Figure S15A). The genotypic groups under moderate drought stress were similar to control conditions but outliers were observed, such as SprattArcher and UnumliArpa (Figure S15C). The before described clustering of inbreds according to their geographical origin was disrupted under dry down stress (Figure S15B).

In the second step, we calculated the stress induces change index SRI for each inbred as the ratio of adjusted entry means under stress condition divided by those of the control condition. Hence, SRI close to one indicated a lower change due to stress, and the reverse is true for SRI lower (PH, TN, LT, RWC, RC, gH^+^ and Φ_II_) or higher than one (Ecs_t_, vH^+^, NPQ_t_ and Φ_NPQ_). In order to get an overview of response to drought stress, we performed a principal component analysis (PCA) using the SRI of all evaluated parameters. SRI of PSII related parameters were the major contributor of principal component 1 (PC1) and SRI of RWC and morphological parameters were the major contributor to PC2 under DD stress. SRI of Φ_NO_ and pmf-related parameters contributed to both PC1 and PC2 under DD stress (Figure 1A). In contrast, both PSII (except Φ_NO_ contributed only to PC1) and pmf related parameters (except gH^+^ contributed only to PC) contributed to variation explained by PC1 and PC2 under MD (Figure 1B).

**Figure 1:**
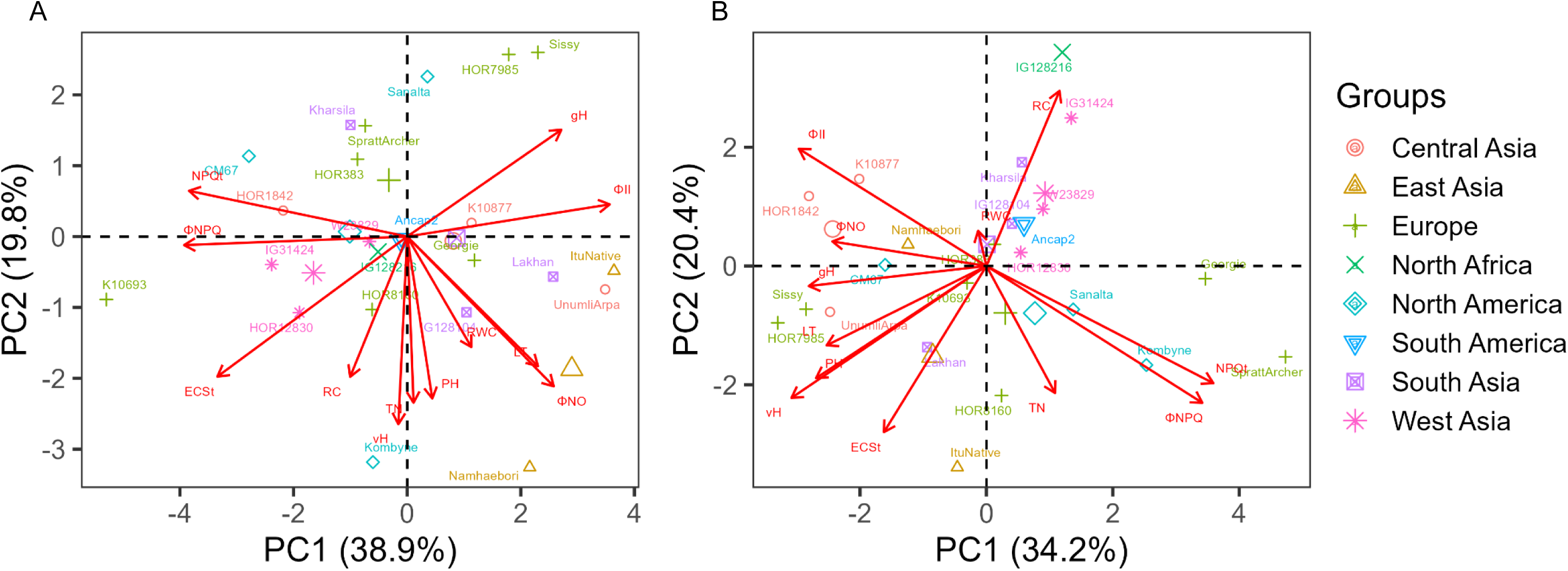
Principle component (PC) analysis of stress response indices (SRI) morphological, physiological and photosynthetic parameters. SRI was calculated as ratio of adjusted entry mean (AEM) of morphological and spectral traits observed in (A) dry down stress versus control and (B) moderate drought versus control to perform principal component analysis. The points represent genotypes and the vectors represent the evaluated parameters namely tiller numbers (TN), plant height (PH), leaf thickness (LT), relative chlorophyll (RC), relative water content (RWC), quantum yield of photosystem II (Φ_II_), non-photochemical losses (Φ_NO_), the fraction of absorbed light dissipated by non-photochemical quenching(Φ_NPQ_ and NPQ_t_), magnitude of electrochromic shift (ECS_t_), proton conductivity (gH^+^) and steady state proton flux (vH^+^). For the dry down treatment, the pots underwent controlled dehydration, and the data were collected for 1 d, 4 d and 7 d after the start of drought stress treatment. For moderate drought stress plants were grown under constant volumetric moisture content of 15% and the data were collected from 1 d, 4 d, 7 d, 11 d, 14 d, 18 d, 21 d, 25 d and 28 d after exposition to MD stress. The experiment was repeated two times with three independent biological replicates per genotype. AEM of the morphological and photosynthetic parameters were estimated across the time points and two independent experiments.

The first two PCs based on SRI also separated the inbreds suggesting that these differ with respect to their drought stress response. For example, PC1 separated K10693 from the rest of the parental inbreds under DD (Figure 1A). K10693 had the highest SRI for NPQ (Φ_NPQ_ and NPQ_t_) and lowest SRI for Φ_II_ (Figure S8, S10 and S11). In contrast, inbreds with low SRI for NPQ (Φ_NPQ_ and NPQ_t_) such as Lakhan, Unumli-Arpa and ItuNative clustered together under DD (Figure 1A). Sissy, HOR7985 and Namhaebori were the other inbreds on the extremes of PC1 and PC2 while Kombyne and Sanalta were on the extremes of PC2 (Figure 1A). Under MD, IG128216 and SprattArcher were the extremes, indicating the strong water-stress response compared to other inbreds. We also observed that the inbreds from South Asia (except Lakhan), West Asia (except IG31424) and those from Central Asia formed distinct clusters (Figure 1B). The best performing inbreds under MD were from Central Asia (HOR1842 and K10877) with high SRI for Φ_II_, Φ_NO_, vH^+^, LT, PH, RC and low SRI for NPQ (Φ_NPQ_ and NPQ_t_) (Figure 1, Figure S7-S14).

We assigned the parental inbreds to drought tolerant and sensitive groups based on the SRI for the evaluated morphological and physiological parameters under MD and DD stress. The inbreds with SRI higher than the median SRI (among the 23 inbreds) of TN, PH, RC, RWC, LT, Φ_II_, Φ_NO_, and gH^+^ were grouped to the tolerant half and the opposite was true for the sensitive half (Table 2). The inbreds with higher SRI compared to median SRI of NPQ (Φ_NPQ_, NPQ_t_), Ecs_t_ and vH^+^ were grouped to the sensitive half and vice versa. We found genotype groups that were sensitive to DD and MD (Group A), (Group B) tolerant to DD and MD, (Group C) tolerant to MD and sensitive or inconsistent response to DD, (Group D) sensitive to MD and tolerant or inconsistent to DD and (Group E) inconsistent response to MD and DD stress (Table 2).

**Table 2:**
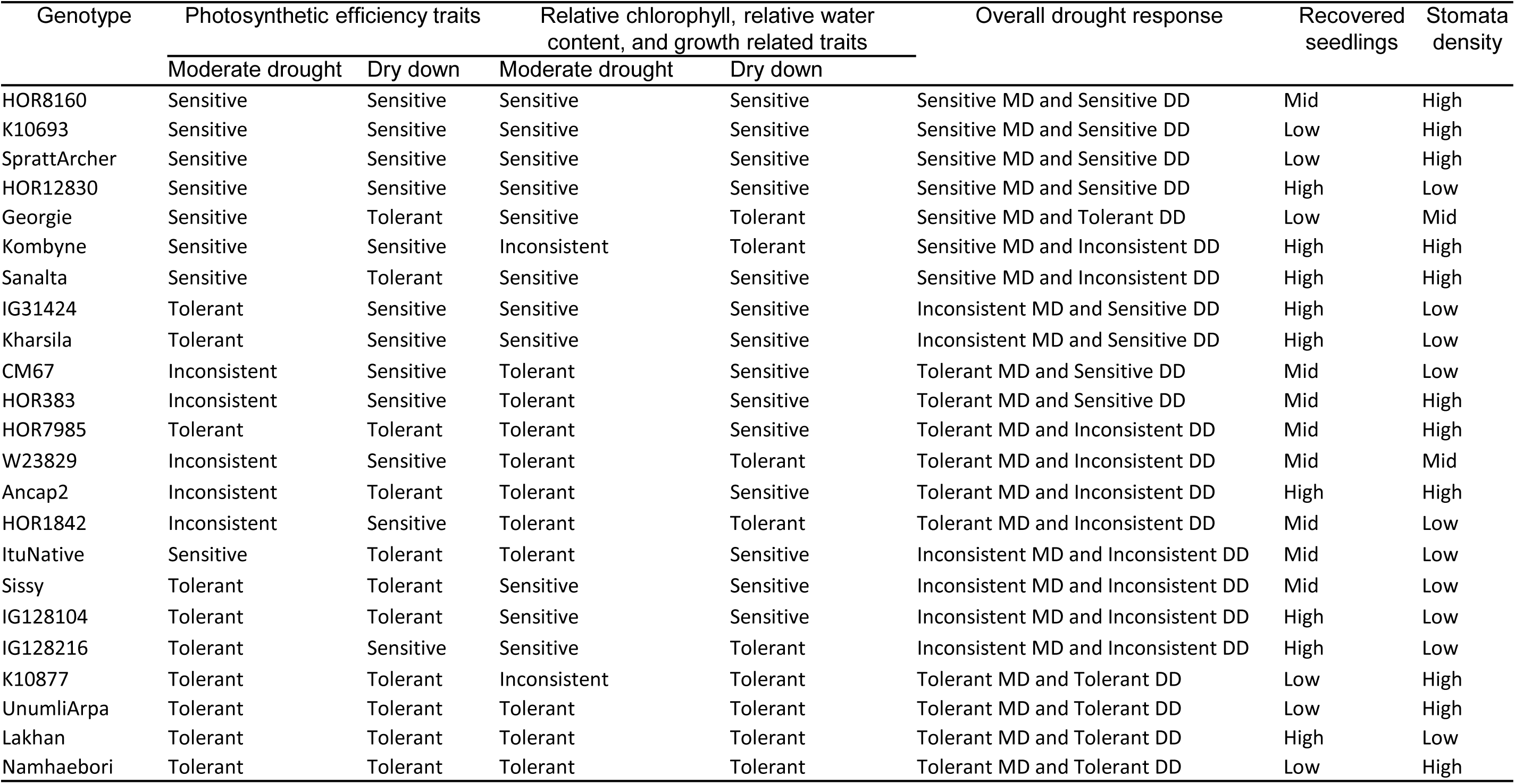
Assignment of genotypes to groups based on the response to dry down (DD) and moderate drought (MD) stress. The inbreds with stress response index (SRI) more than the median SRI (among the 23 inbreds) of tiller number, plant height, relative chlorophyll index, relative water content, leaf thickness, quantum yield of photosystem II (Φ_II_), non-photochemical losses (Φ_NO_), and proton conductivity (gH^+^) were grouped to tolerant half and opposite was true for sensitive half. The inbreds with higher SRI compared to median SRI of NPQ (Φ_NPQ_ and NPQ_t_), magnitude of electrochromic shift (ECS_t_) and steady state proton flux (vH^+^) were grouped to sensitive half and vice versa.

| Genotype | Photosynthetic efficiency traits |  | Relative chlorophyll, relative water content, and growth related traits |  | Overall drought response | Recovered seedlings | Stomata density |
| --- | --- | --- | --- | --- | --- | --- | --- |
|  | Moderate drought | Dry down | Moderate drought | Dry down |  |  |  |
| HOR8160 | Sensitive | Sensitive | Sensitive | Sensitive | Sensitive MD and Sensitive DD | Mid | High |
| K10693 | Sensitive | Sensitive | Sensitive | Sensitive | Sensitive MD and Sensitive DD | Low | High |
| SprattArcher | Sensitive | Sensitive | Sensitive | Sensitive | Sensitive MD and Sensitive DD | Low | High |
| HOR12830 | Sensitive | Sensitive | Sensitive | Sensitive | Sensitive MD and Sensitive DD | High | Low |
| Georgie | Sensitive | Tolerant | Sensitive | Tolerant | Sensitive MD and Tolerant DD | Low | Mid |
| Kombyne | Sensitive | Sensitive | Inconsistent | Tolerant | Sensitive MD and Inconsistent DD | High | High |
| Sanalta | Sensitive | Tolerant | Sensitive | Sensitive | Sensitive MD and Inconsistent DD | High | High |
| IG31424 | Tolerant | Sensitive | Sensitive | Sensitive | Inconsistent MD and Sensitive DD | High | Low |
| Kharsila | Tolerant | Sensitive | Sensitive | Sensitive | Inconsistent MD and Sensitive DD | High | Low |
| CM67 | Inconsistent | Sensitive | Tolerant | Sensitive | Tolerant MD and Sensitive DD | Mid | Low |
| HOR383 | Inconsistent | Sensitive | Tolerant | Sensitive | Tolerant MD and Sensitive DD | Mid | High |
| HOR7985 | Tolerant | Tolerant | Tolerant | Sensitive | Tolerant MD and Inconsistent DD | Mid | High |
| W23829 | Inconsistent | Sensitive | Tolerant | Tolerant | Tolerant MD and Inconsistent DD | Mid | Mid |
| Ancap2 | Inconsistent | Tolerant | Tolerant | Sensitive | Tolerant MD and Inconsistent DD | High | High |
| HOR1842 | Inconsistent | Sensitive | Tolerant | Tolerant | Tolerant MD and Inconsistent DD | Mid | Low |
| ItuNative | Sensitive | Tolerant | Tolerant | Sensitive | Inconsistent MD and Inconsistent DD | Mid | Low |
| Sissy | Tolerant | Tolerant | Sensitive | Sensitive | Inconsistent MD and Inconsistent DD | Mid | Low |
| IG128104 | Tolerant | Tolerant | Sensitive | Sensitive | Inconsistent MD and Inconsistent DD | High | Low |
| IG128216 | Tolerant | Sensitive | Sensitive | Tolerant | Inconsistent MD and Inconsistent DD | High | Low |
| K10877 | Tolerant | Tolerant | Inconsistent | Tolerant | Tolerant MD and Tolerant DD | Low | High |
| UnumliArpa | Tolerant | Tolerant | Tolerant | Tolerant | Tolerant MD and Tolerant DD | Low | High |
| Lakhan | Tolerant | Tolerant | Tolerant | Tolerant | Tolerant MD and Tolerant DD | High | Low |
| Namhaebori | Tolerant | Tolerant | Tolerant | Tolerant | Tolerant MD and Tolerant DD | Low | High |

### Metabolite profile under drought stress

We profiled metabolites under all three environmental conditions using an untargeted approach. The sample collection for metabolite profiling was done at a single day with only one reference control group which was different to the before described morphological and physiological parameters. From this experiment, we obtained the relative quantification of 42 metabolites that could be broadly classified into amino acids, low molecular weight organic acids, sugars, polyhydroxy acids and fatty acid derivatives (Figure 2). We observed significant (p < 0.05-0.001) genotypic differences for all the metabolites. Significant (p < 0.05-0.001) treatment and genotype by treatment interaction effects were detected for almost all metabolites (Table S4). For more than two-thirds of the metabolites, heritability was higher than 47 % (Control), 37 % (DD) and 50 % (MD) (Table S5).

**Figure 2:**
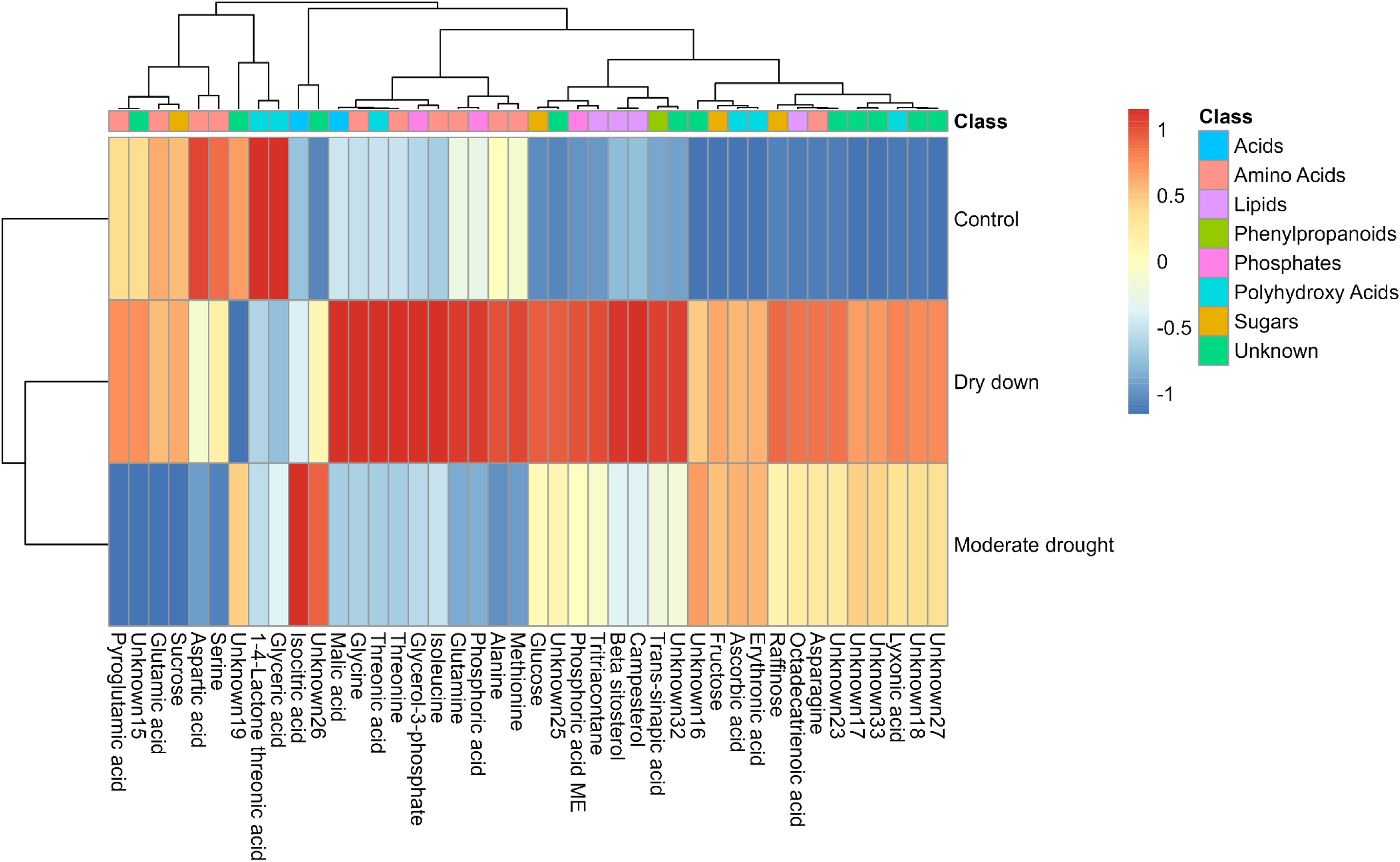
Heatmap of the adjusted entry mean of 42 metabolites detected in all 23 genotypes under control condition and two types of drought stress. Dark red and blue colors indicate the higher and lower relative value of the metabolite, respectively. The metabolite profiling was done in the leaf samples collected from 7 d and 12 d after stress under dry down and moderate drought stress, respectively. The leaves from the control conditions collected at 7 d and 12 d were pooled together making one representative control group. The experiment was repeated two times with three independent biological replicates per genotype. Adjusted entry mean of the metabolites were estimated across two independent experiments.

The principal component analysis based on the data from the metabolite profiling separated the metabolite accumulation pattern in control conditions from water stress (DD and MD). The DD and MD indicated overlapping as well as distinct responses for seven different types of primary metabolites (Figure 2). Amino acids and low molecular weight organic acids were highly abundant under DD compared to MD and control conditions. Except for sucrose, the sugars such as glucose, fructose and raffinose showed higher accumulation under DD and MD compared to control conditions. The majority of the unknown metabolites (six out of nine) behaved similar to sugars with higher abundance under DD and MD stress compared to control conditions. In contrast, fatty acid derivatives were highly abundant under DD, followed by MD and less abundant under control conditions (Figure 2).

Next, we constructed the heat maps of SRI for DD and MD for those metabolites that showed higher relative quantities compared to control conditions. Under DD, the dendrogram can be broadly broken into four different groups (Figure 3A). The first one was Lakhan type, that showed a higher abundance of monosaccharides (glucose and fructose), ascorbic acid and unknown33 compared to the rest of the inbreds. The second group was UnumliArpa type, where higher SRI for all metabolites under DD was observed compared to other groups of inbreds. W23829/8011 was the third group that had higher SRI for other metabolites except glucose and fructose (Figure 3A). A large group of 10 inbreds with low-to-moderate SRI for all metabolites formed the fourth group (Figure 3A). Next, we clustered the inbreds according to the metabolites with higher relative values under MD compared to control conditions. Lakhan, UnumliArpa and IG128216 behaved similarly with higher SRI for sugars, ascorbic acid, polyhydroxy acids and six unknown metabolites than other inbreds. Another group of seven inbreds showed low SRI for the above-mentioned metabolites (Figure 3B).

**Figure 3:**
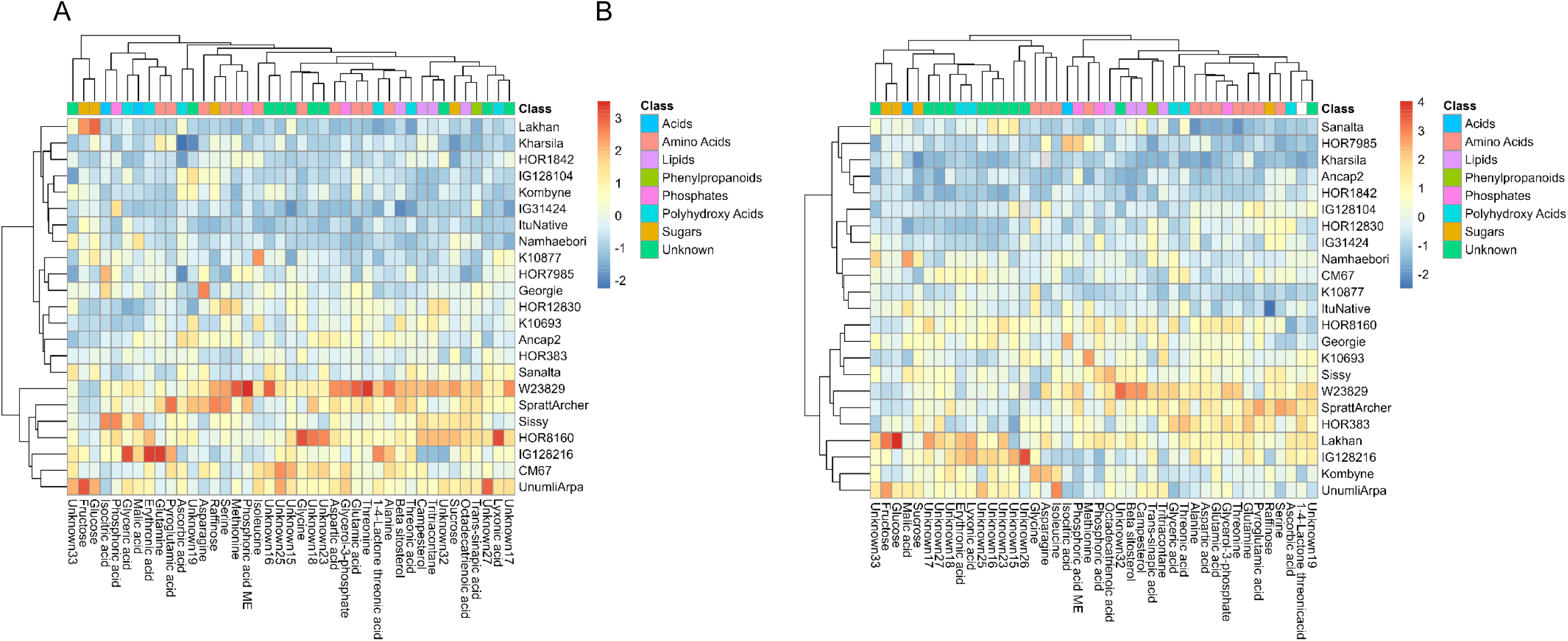
Heatmap of stress response index (SRI) of 42 metabolites described in Figure 2 across 23 barley inbreds. SRI was calculated as ratio of adjusted entry means (AEM) of metabolites accumulated in (A) dry down (DD) stress versus control condition (B) moderate drought (MD) stress versus control condition. Dark red and blue colors indicate the higher and lower SRI of the metabolites, respectively. The metabolite profiling was done in the leaf samples collected from 7 d and 12 d after the start of DD and MD stress treatment, respectively. The leaves from the control conditions collected at 7 d and 12 d were pooled together making one representative control group. The experiment was repeated two times with three independent biological replicates per genotype. AEM of the metablotes were estimated across two independent experiments.

### Variation of stomata density among barley inbreds

We measured the SD from the adaxial and abaxial surface of the leaf from the bottom, middle and upper third of the leaf. We observed significant (p < 0.001) variation in SD among the inbreds, leaf surface and the position on the leaf blade as well as significant inbred by position interaction effect (Table S2). The *h^2^* of SD was 77% to 88% at different leaf positions. SD ranged from 18 to 33, 21 to 34 and 28 to 38 stomata/mm^2^ on the bottom, middle and top positions of the leaf blade, respectively. The PCA of the SD information indicated that the SD are similar in the middle and the bottom parts on the leaf blade but different from the SD towards the tip of the leaf blade (Figure 4). PC1 separated the inbreds from Europe with those from Asia and Africa, while PC2 separated the inbreds from European origin (Figure 4). HOR1842 and HOR7985 were the inbreds with highest and lowest SD, respectively (Figure 5C).

**Figure 4:**
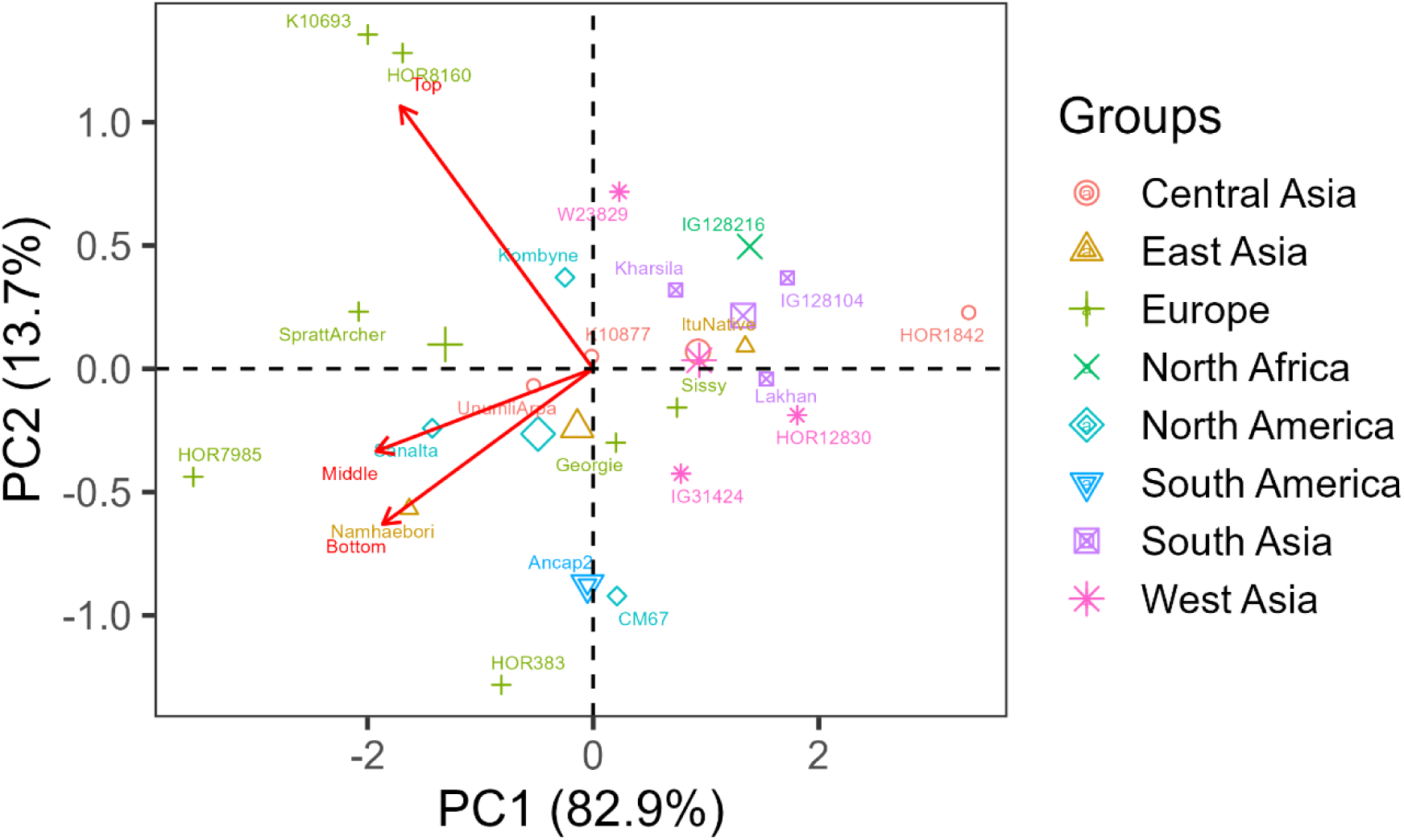
Principle component (PC) analysis of adjusted entry mean of stomata density. Stomata density was measured on the abaxial (lower surface) and the adaxial (upper surface) of the leaves at three different position on the leaf blade. Epidermal imprints were taken from two-week-old seedlings to estimate stomata density. The experiment was repeated two times with three independent biological replicates per genotype. AEM of stomata density were estimated across two independent experiments and leaf surfaces at three different positions on the leaf blade.

**Figure 5:**
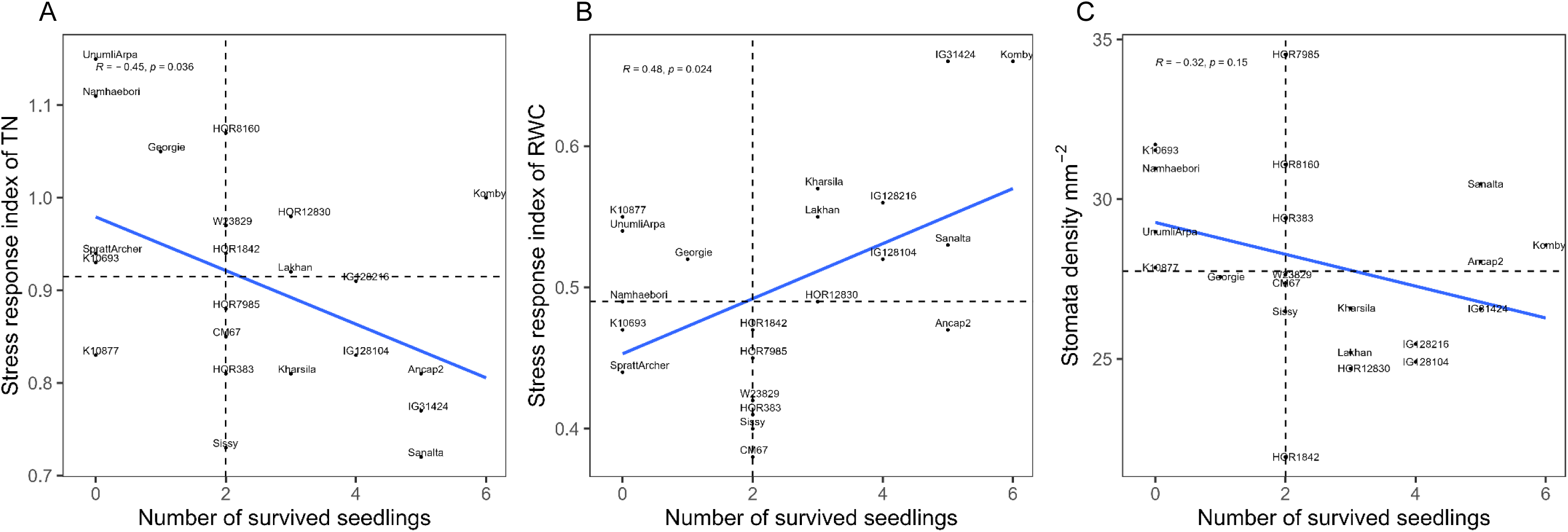
Correlation of the count of recovered plants and morphological and physiological parameters. Stress response index calculated as the ratio of adjusted entry mean (AEM) for (A) tiller number (TN) and (B) relative water content observed in dry down stress versus control conditions; and (C) AEM of stomata density were used to estimate linear relationship with count of recovered plants. For the dry down stress, the pots underwent controlled dehydration, and the data were collected for 1 d, 4 d and 7 d after the start of drought stress treatment. The experiment was repeated two times with three independent biological replicates per genotype. AEM was estimated across the time points and two independent experiments. Plants were rewatered 12 d after the start of dry down stress. Then, the plants that produced true leaves and resumed growth two weeks after the start of rewatering were counted as recovered plants. AEM of stomata density were estimated across two independent experiments at three different leaf position from two abaxial and adaxial surface of leaf blade. The vertical and horizontal dashed lines indicate the median value of the x-axis and y-axis variables, respectively. The diagonal blue line is the trend-line of the linear relationship between the variables.

### Correlation of morpho-physiological and metabolic adjustments with the recovery of plants after dehydration stress

The watering was discontinued in plants under DD until 12 d after the start of stress. Afterwards, the plants were rewatered and phenotypically characterized. We counted for each inbred the number of plants that were able to produce true leaves 14 d after rewatering. All plants of Kombyne recovered, while for the inbreds K10693, K10877, Namhaebori, SprattArcher and UnumliArpa no plants recovered (Figure 5). The count of recovery was negatively correlated with SRI of TN (R = −0.45, p < 0.05) and positively correlated with SRI of RWC under DD stress (R = 0.48, p < 0.05) (Figure 5A and B). Another morphological feature among the inbreds with low recovery count was the high SD (Figure 5C). For instance, the SD of the inbreds with low survival count was significantly higher (p < 0.05) than those with high recovery (Figure S16).

## Discussion

In our study, we combined the comparative analysis of plant morphology, photosynthetic efficiency and metabolite abundance at varying soil moisture status in a set of barley inbreds selected to maximize phenotypic and genotypic diversity (Weisweiler *et al*., 2019). In addition, these 23 inbreds which comprised landraces and cultivars with worldwide origin are also the parents of the HvDRR population (Casale *et al*., 2022).

### Barley uses different strategies to tolerate moderate drought and dry down stress

Heritability of PSII and pmf related parameters was moderate to high across the two experimental repetitions (Table S3). Our study represents only the second instance of using MutispeQ for large-scale phenotyping of diverse genotypes in barley under drought stress. The only previous study on barley that used MutispeQ (Fernández-Calleja *et al*., 2020) did not report repeatability of the PSII (and the pmf related parameters were not included) related parameters for meaningful comparison. Nevertheless, highly repeatable observations of heritability of PSII and pmf related parameters across different growing conditions and repeated experiments in the current study showed the reliability and robustness of MultispeQ for genetic screening of barley germplasm under drought stress and beyond.

Likewise, the intensity of drought stress affected the heritability of metabolites, as it considerably decreased under DD stress compared to MD stress and control conditions (Table S5). The *h^2^* of plant morphological parameters was high and it decreased only marginally under MD and DD stress compared to control conditions (Table S3). Therefore, changes in the PSII and pmf related parameters, primary metabolite profile, and plant morphology related parameters could therefore be used as selection parameters for different levels of drought tolerance in barley.

Starting with PSII related parameters, we observed a significant decrease in Φ_II_ and Φ_NO_ (SRI < 1) as well as an increase in Φ_NPQ_ (SRI > 1) under DD and MD stress vs control conditions (Figure S8-S11). This was expected because the capacity of light absorption to utilization might be lower in plants under drought stress than control conditions. This is due to the reduced CO_2_ exchange coupled with damages of the photosynthetic apparatus (Pinheiro and Chaves, 2011). Therefore, NPQ is activated in plants under drought to prevent photo-inhibition (Kanazawa and Kramer, 2002*a*). Hence, the protective mechanism (NPQ) competes with the utilization of absorbed light energy, causing reduced quantum yield of photosystem II (Müller *et al*., 2001). Although Φ_II_ declines with drought stress in our study, some studies reported an increase in Φ_II_ especially under MD stress (Wang *et al*., 2018*b*; Yao *et al*., 2018). In contrast, SRI of Φ_II_ did not differ in our study between DD and MD when averaged across all genotypes. However, SRI of NPQ was 7.6% higher while that of Φ_NO_ was lower (5%) in MD compare to DD (Figure S9-S11). Elevated values of Φ_NO_ indicate that a plant cannot defend itself against the harmful effects of excessive light (Kramer *et al*., 2004; Klughammer and Schreiber, 2008). Therefore, the photo-protective capacity to compensate for excess illumination is more efficient in plants under MD compared to DD stress. Averaged across all inbreds, we observed that the change in PSII related parameters under DD vs control conditions and the MD vs control conditions were not the same, which indicated the different strategies to tolerate MD and DD stress.

Here, we reported for the first instance the changes in the total pmf (ECS_t_), thylakoid conductance (gH^+^) and proton flux (vH^+^) under drought stress in barley. Averaged across 23 inbreds, gH^+^ decreased in barley under DD and MD stress compared to control conditions (Figure S13). Low CO_2_ levels caused a decrease in gH^+^ non-stressed plants of Arabidopsis and tobacco by decreasing ATP synthase activity (Kanazawa and Kramer, 2002*b*; Avenson *et al*., 2005*b*). Therefore, low CO_2_ exchange due to drought induced stomata closure might have caused lower gH^+^ under drought stress than well-watered conditions. It was previously reported that the stomatal conductance and intracellular CO_2_ concentration decreased proportionally with drought intensity in barley (Shrestha *et al*., 2022). Because the reduction of gH^+^ was more pronounced in MD (10% lower) than in DD stress, CO_2_ exchange alone cannot account for the change in gH^+^, especially under drought stress. Alternatively, the proton leak across the thylakoid membrane in severe drought stress might account for higher gH^+^ under DD compared to MD stress. Next, compared to control plants, we observed a rise in the ECS_t_ and vH^+^ under DD stress compared to MD stress (Figure S13-S14). The change in CO_2_ levels had no effect on total pmf (ECS_t_) and vH^+^ in non-stressed Arabidopsis plants (Avenson *et al*., 2005*a*,*b*). However, many studies have well-documented that elevation of cyclic electron flow under drought stress as the lumen acidification limits linear electron flow (Zivcak *et al*., 2014). Kohzuma *et al*. (2009) found that cyclic electron flow was the major contributor of net increase in ECS_t_ and vH^+^ under drought stress in melon. Although the change in ECS_t_ was similar in DD and MD stress compared to control conditions (Figure S12), SRI of vH^+^ was 12% higher in DD compared to MD in barley (Figure S14). Therefore, the net increase in pmf alone did not fully explain the difference in vH^+^ between DD and MD stress (Figure S14). As stated earlier in the case of gH^+^, the differences in vH^+^ might be caused by the introduction of oxidative damage to the thylakoid membrane. Because of the higher water loss under DD stress (42% RWC) compared to MD stress (71% RWC) contribute to greater oxidative damage to the chloroplast component in DD stress than in MD stress (Figure S3). This was also clear from the RC, which was stable under MD (no treatment effect) while the RC values decreased over time under DD (Figure S7).

The change in the metabolite profile also differed between the growing conditions and the intensity of drought stress (Figure 2 and 3). The pooled control samples formed a distinct cluster in terms of metabolite profile compared to DD (sampling at 7 d after stress treatment) and MD (sampling at 12 d after stress treatment) stress. The pooling of samples from control conditions should not introduce significant bias when used as a single reference in comparisons to the metabolite from those of DD and MD stress. This is because the difference in metabolite profile in the same inbred within 5 d under the same growing conditions is unlikely. This is in agreement with previous experiments in wheat (Bowne *et al*., 2012*a*) and barley (Piasecka *et al*., 2017) that revealed a minimal shift or high correlation in metabolite profile in leaf over a 3 d to 15 d period under well-watered conditions.

Overall, the metabolic activity was higher in plants under drought stress and it increased with the intensity of drought stress (Figure 2 and 3). Consistent with the literature, we observed a significant up-regulation of the primary metabolite group such as sugars, amino acids, polyhydroxy acids, polyols and fatty acids under drought stress compared to control conditions (Templer *et al*., 2017; Lawas *et al*., 2019; Itam *et al*., 2020). We discovered that hexoses and polyhydroxy acid accumulated at similar levels under MD and DD stress, while lipids, phosphates and amino acids were more upregulated under DD stress compared to MD and control conditions. The change in the metabolite pool concurred with previous observations in barley, wheat, rice and other plant species (Todaka *et al*., 2017*b*; Fàbregas and Fernie, 2019).

Compared to control conditions, reduction in shoot growth parameters was more pronounced in MD than in DD stress (Figure S4-S6). Plants have more time to adapt under MD stress and, thus, the growth retardation might result as the effect of reduced photosynthetic capacity (Sakoda *et al*., 2022) as well as feedback response to decrease the size of photosynthetic organs which are responsible for transpirational loss which in turn alters the root to shoot ratio (Xu *et al*., 2015). Hence, the shoot growth will continue to grow at reduced capacity as an adaptive response, adjusting to available soil moisture (Skirycz and Inzé, 2010; Claeys and Inzé, 2013). In contrast, due to immediate and acute nature of DD stress, plant apply drastic physiological changes to survive stress and the timeframe to activate adaptive strategies is limited compared to MD stress.

The above described differences in stress response between MD and DD stress are an average reaction across the investigated inbreds. However, we observed significant genotype-dependent reactions to DD and MD stress, which is discussed in the following section.

### Drought tolerant parental inbreds of HvDRR population

We assigned the parental inbreds to drought tolerant and sensitive groups based on the SRI for the evaluated morphological and physiological parameters under moderate drought and dry down stress (Table 2). We found five distinct groups of genotypes. Except HOR12830, all inbreds sensitive to DD and MD originated in Europe, while the inbreds tolerant to DD and MD originated in East, South and Central Asia (Table S1 and Table 2). The barley accessions domesticated in semi-arid sub-tropical regions in South and Central Asia might have accumulated genetic and physiological features to tolerate drought stress (Elakhdar *et al*., 2022). Inbreds that were only tolerant to MD (Group C) predominantly came from the temperate regions in Europe and Americas (Table 2 and Table S1). Historically, barley cultivated and bred in the temperate region were exposed to favorable soil-water regime and water stress in the region are usually mild (Templer *et al*., 2017). Hence, it is not surprising to observe that the genotypes originating from this region are either sensitive to drought stress or tolerate only mild stress.

Next, as a proxy for drought resistance, we exposed plants to severe DD stress for 12 days and counted the plants that produced true leaves after rewatering. The plants that had high survival count used conservative strategies under drought stress and performed poorly for physiological and morphological adjustments under DD and MD stress (Figure 5). Hence, consistent with previous reports, our study also confirmed trade-offs between survival and growth performance under drought stress (Claeys and Inzé, 2013). Another common feature among the inbreds with low recovery count was low RWC under DD and high SD (Figure 5, Figure S16). The major source of water loss from the plants occurs through stomata. High SD are often associated with reduced water use efficiency and poor drought survival (Hughes *et al*., 2017). Kombyne and Ancap2 were the only exceptions with high SD and a very high recovery count. Hence, the altered stomata regulation by hormonal signaling might also contribute to conserve leaf tissue water content in Kombyne and Ancap2 which requires further research.

The results of our study suggested that some barley inbreds endured drought by continuing plant growth and efficient photosynthesis, while a few inbreds use a survival strategy by conserving tissue hydration and, investing in protective processes and reduced growth under drought. For instance, low SRI of tiller numbers and high RWC were the common features of the latter inbreds (Figure 5A and Figure S16). Therefore, the genotypes that can restrict growth during stress might have higher chances of surviving prolonged drought. Therefore, the HvDRR population, which was derived from pair-wise crosses among the studied inbred lines, is a useful resource for mapping the genetic control of intra-species differences in metabolite response and drought tolerance strategy in barley.

### Association between metabolic activity and drought stress response

Several *in vitro* and *in vivo* studies have claimed that the increase in primary metabolite pool, such as soluble sugars and amino acids, are essential for abiotic stress tolerance, including drought stress (Graham and Wilkinson, 1992; Hare *et al*., 1998; Székely *et al*., 2008; Sharma *et al*., 2011; Merewitz *et al*., 2012; Muzammil *et al*., 2018; Ashrafi *et al*., 2018; Valitova *et al*., 2019). We compared the metabolite profile of the inbreds to understand the relationship between metabolic activity and performance under drought stress. We referred to the inbreds with high metabolic activity as the ones that showed higher SRI, which was measured as the ratio of AEM of relative metabolite value under drought stress (DD or MD) versus control conditions (Figure 1 and 2, Figure S17-S20). The difference in the metabolism of hexose (glucose and fructose) and metabolites of unknown type sets apart the inbreds of Group B (tolerant to both DD and MD stress) from the rest. Especially UnumliArpa was the most metabolically active inbred under MD and DD, respectively (Figure 2 and 3). In support of such metabolic differences between sensitive and tolerant inbreds, it has been demonstrated that simple sugars accumulated under MD helps to maintain plant growth and those compartmentalized in vacuole offers a first line of defense against water loss as reported for barley, wheat and Arabidopsis (Fàbregas and Fernie, 2019).

Three sensitive inbreds SprattArcher, HOR8160 (group A, sensitive to DD and MD) and W23829 (group C, sensitive to DD) were among the highest accumulators of sucrose, amino acids, lipids and organic acid under MD and DD (Figure 3) indicating that such type of metabolites might accumulate to counter the negative effect of rapid water loss (Bowne *et al*., 2012*b*; You *et al*., 2019; Djemal and Khoudi, 2021). This was in agreement with low RWC in those inbreds. In addition to the above observations, lower metabolism of primary metabolites especially under DD stress was the major metabolic response of inbreds that showed high survival under severe dehydration. For instance, five inbreds with high recovery count formed a distinct cluster with lower metabolic activity under DD stress. IG128216 was the only inbred that had higher metabolic activity under MD and DD stress (Figure 3). Therefore, the genotypes with better drought resistance were also metabolically less active compared to the rest and followed conservative approach under DD stress. These observations suggest that (i) early sugar metabolism might have more positive impact on drought tolerance than other groups of primary metabolites and (ii) accumulation of metabolites alone cannot determine drought tolerance level and more is better concept should be used with caution.

## Acknowledgement

The authors would like to thank the greenhouse staff of the Heinrich-Heine University Düsseldorf for the logistics to carryout experiments in the greenhouse. Additionally, we express our thanks to Gadi Vogeler for assisting with leaf sampling and measuring soil volumetric moisture content. We also acknowledge the computational platform provided by the Center of Information and Media Technology at Heinrich-Heine-University Düsseldorf.

## Authors contribution

Conceptualization of the study: AS and BS

Experimentation: AS, TK and LAM

Metabolite profiling: PW and AE

Data processing: AS and BS

Manuscript preparation: AS and BS

Supervision: AS and BS

Funding acquisition: BS

All authors have read and approved the manuscript.

---

**Table S1:** List of genotypes used in the study.

| Genotype | Gene bank ID | Country | Continent | Germplasm type | Row type |
| --- | --- | --- | --- | --- | --- |
| Ancap2 | BCC807 | URY | South America | cultivar | 6 |
| CM67 | BCC846 | USA | North America | cultivar | 6 |
| Georgie | BCC1381 | GBR | Europe | cultivar | 2 |
| HOR12830 | HOR12830 | SYR | Africa | landrace | 6 |
| HOR1842 | HOR1842 | AFG | Asia | landrace | 6 |
| HOR383 | BCC1561 | BGR | Europe | landrace | 6 |
| HOR7985 | HOR7985 | TUR | Europe | landrace | 2 |
| HOR8160 | HOR8160 | TUR | Europe | landrace | 2 |
| IG128104 | BCC173 | PAK | Asia | landrace | 6 |
| IG128216 | BCC118 | LBY | Africa | landrace | 6 |
| IG31424 | BCC190 | SYR | Africa | landrace | 2 |
| ItuNative | BCC502 | CHN | Asia | cultivar | 6 |
| K10693 | BCC1491 | RUS | Europe | landrace | 6 |
| K10877 | BCC1503 | TKM | Asia | landrace | 6 |
| Kharsila | HOR11403 | IND | Asia | landrace | 6 |
| Kombyne | BCC893 | USA | North America | cultivar | 6 |
| Lakhan | BCC533 | IND | Asia | cultivar | 6 |
| Namhaebori | BCC667 | KOR | Asia | cultivar | 6 |
| Sanalta | BCC929 | CAN | North America | cultivar | 2 |
| Sissy | BCC1413 | GER | Europe | cultivar | 2 |
| SprattArcher | BCC1415 | GBR | Europe | cultivar | 2 |
| UnumliArpa | BCC1470 | UZB | Asia | cultivar | 2 |
| W23829/803911 | HOR11374 | ISR | Asia | landrace | 2 |

**Table S2:** Significance of experimental factors and their interactions on the variation of stomata density. Asterisks indicate the level of significance (***, p ≤ 0.001; ns, p > 0.05). The experimental factors include genotype, two independent experiments (Rep), leaf surface and position on the leaf blade (Position). Stomata density was measured on the abaxial (lower surface) and the adaxial (upper surface) of the leaves at three different position on the leaf blade. Epidermal imprints were taken from two-week-old seedlings to estimate stomata density. The experiment was repeated twice with three independent biological replicates per genotype. Heritability (*H^2^*) of stomata density were estimated across two independent experiments and leaf surfaces at three different positions on the leaf blade.

| Parameters | Significance level and $H^2$ |
| --- | --- |
| Genotype | *** |
| Rep | *** |
| Surface | *** |
| Position | *** |
| Genotype:Surface | ns |
| Genotype:Position | *** |
| $H^2$ (Top) | 0.77 |
| $H^2$ (Middle) | 0.8 |
| $H^2$ (Bottom) | 0.88 |

**Table S3:** Broad sense heritability of morphological, physiological and photosynthetic parameters under control and stress conditions. For the dry down treatment, the pots underwent controlled dehydration, and the data were collected for 1 d, 4 d and 7 d after the start of drought stress treatment. For moderate drought stress plants were grown under constant volumetric moisture content of 15% and the data were collected from 1 d, 4 d, 7 d, 11 d, 14 d, 18 d, 21 d, 25 d and 28 d after exposition to MD stress. The experiment was repeated two times with three independent biological replicates per genotype. Heritability of the evaluated parameters were estimated across the time points and two independent experiments. Abbreviations: Non-photochemical quenching of chlorophyll fluorescence in photosystem II (Φ_NPQ_ and NPQ_t_), magnitude of electrochromic shift (ECS_t_), conductivity (gH^+^), steady state proton flux (vH^+^).

| Stress type | Trait | Control | Stress |
| --- | --- | --- | --- |
| Dry down | $\Phi_{II}$ | 0.42 | 0.69 |
| | $\Phi_{NO}$ | 0.59 | 0.52 |
| | $\Phi_{NPQ}$ | 0.15 | 0.65 |
| | $NPQ_t$ | 0.24 | 0.59 |
| | $ECS_t$ | 0.80 | 0.86 |
| | $gH^+$ | 0.63 | 0.70 |
| | $vH^+$ | 0.67 | 0.82 |
|  | Relative water content (%) | 0.34 | 0.31 |
|  | Relative chlorophyll index | 0.84 | 0.80 |
| | Leaf thickness ($\mu m$) | 0.94 | 0.85 |
|  | Plant height (cm) | 0.95 | 0.92 |
|  | Tiller number | 0.79 | 0.85 |
| Moderate drought | $\Phi_{II}$ | 0.61 | 0.71 |
| | $\Phi_{NO}$ | 0.78 | 0.72 |
| | $\Phi_{NPQ}$ | 0.55 | 0.69 |
| | $NPQ_t$ | 0.59 | 0.67 |
| | $ECS_t$ | 0.91 | 0.84 |
| | $gH^+$ | 0.42 | 0.61 |
| | $vH^+$ | 0.90 | 0.88 |
|  | Relative water content (%) | 0.13 | 0.58 |
|  | Relative chlorophyll index | 0.94 | 0.88 |
| | Leaf thickness ($\mu m$) | 0.96 | 0.95 |
|  | Plant height (cm) | 0.98 | 0.99 |
|  | Tiller number | 0.95 | 0.95 |

**Table S4:** Significance of experimental factors and their interactions on the variation of metabolite profile. The experimental factors include genotype, treatment, two independent experiments (Rep) and tray used to organize the pots. Asterisks indicate the level of significance (***, p ≤ 0.001; **, p ≤ 0.01; *, p ≤ 0.05; ns, p > 0.05). The metabolite profiling was done in the leaf samples collected from 7 d and 12 d after the start of dry down and moderate drought stress, respectively. The leaves from the control conditions collected at 7 d and 12 d were pooled together making one representative control group. The experiment was repeated two times with three independent biological replicates per genotype.

| Trait | G | T | G:T | T:Rep | T:Rep:Tray |
| --- | --- | --- | --- | --- | --- |
| Alanine | *** | *** | ** | *** | *** |
| Asparagine | * | *** | ns | *** | ns |
| Aspartic acid | *** | *** | ** | *** | ns |
| Beta-sitosterol | *** | *** | ns | *** | *** |
| Campesterol | *** | *** | * | * | ** |
| Glutamic acid | *** | *** | ns | *** | * |
| Glutamine | *** | *** | ns | *** | *** |
| Glycine | * | *** | ns | *** | *** |
| Isocitric acid | *** | *** | ** | *** | *** |
| log10(Isoleucine) | * | *** | ns | *** | * |
| Malic acid | *** | *** | *** | *** | *** |
| Methionine | ** | *** | ns | *** | ns |
| Octadecatrienoic acid | *** | ** | ** | *** | *** |
| Pyroglutamic acid_Glutamine_Glutamic acid | *** | *** | ** | *** | *** |
| Serine | ** | *** | * | *** | * |
| Threonine | *** | *** | ** | *** | *** |
| Tritriacontane | *** | *** | ** | ** | *** |
| Unknown15 | *** | *** | * | *** | *** |
| Unknown16 | *** | *** | * | ns | * |
| Unknown33 | *** | *** | * | ns | *** |
| Unknown32 | *** | *** | * | *** | *** |
| Unknown27 | *** | *** | * | *** | *** |
| Unknown26 | *** | *** | ** | *** | ns |
| Unknown25 | *** | *** | * | ** | * |
| Unknown23 | *** | *** | * | ** | ** |
| Unknown19 | *** | ns | ns | * | *** |
| Unknown18 | *** | *** | ** | *** | ns |
| Unknown17 | *** | *** | ** | *** | ns |
| log10(trans-sinapic acid) | *** | *** | ns | * | *** |
| Threonic acid | *** | *** | ns | ns | *** |
| Sucrose | *** | ** | ns | ns | ** |
| Raffinose | *** | *** | *** | ** | *** |
| Phosphoric acid | *** | *** | ** | *** | *** |
| Phosphoric acid monomethyl ester | *** | *** | ** | ns | *** |
| Lyxonic acid | *** | *** | ** | *** | ns |
| Glycerol-3-phosphate | *** | *** | * | *** | *** |
| Glyceric acid | *** | *** | ** | *** | *** |
| 1,4-Lactone threonic acid | *** | ns | ns | ns | *** |
| Ascorbic acid | ** | ns | * | ** | ** |
| Erythronic acid | *** | *** | ns | ** | ** |
| Fructose | *** | *** | *** | *** | *** |
| Glucose | *** | *** | *** | *** | *** |

**Table S5:**
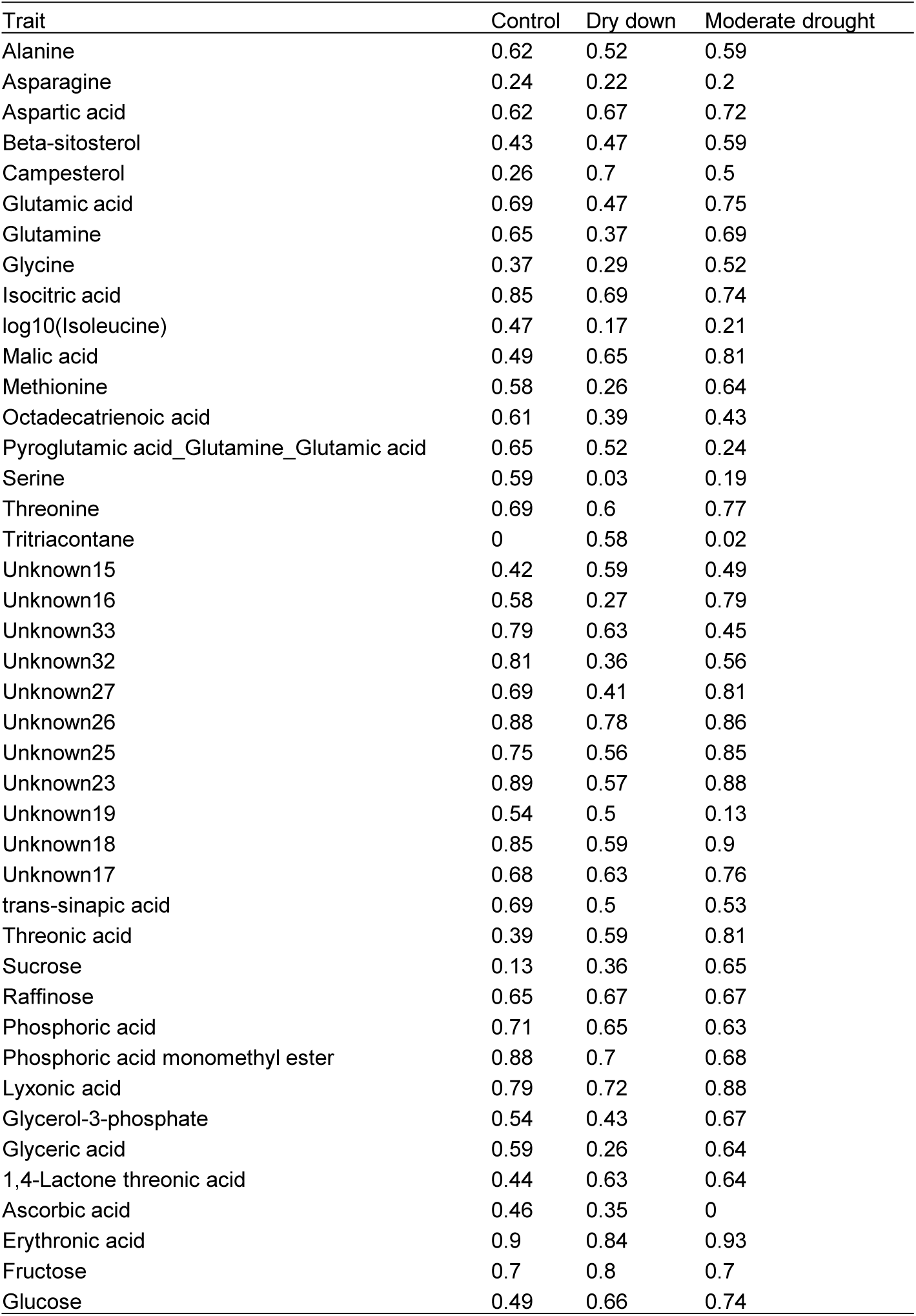
Broad sense heritability of metabolite profile under control, dry down and moderate drought stress. The metabolite profiling was done in the leaf samples collected from 7 d and 12 d after the start of dry down and moderate drought stress, respectively. The leaves from the control conditions collected at 7 d and 12 d were pooled together making one representative control group. The experiment was repeated two times with three independent biological replicates per genotype. Heritability of the metabolite profile was estimated across two independent experiments.

**Figure S1:**
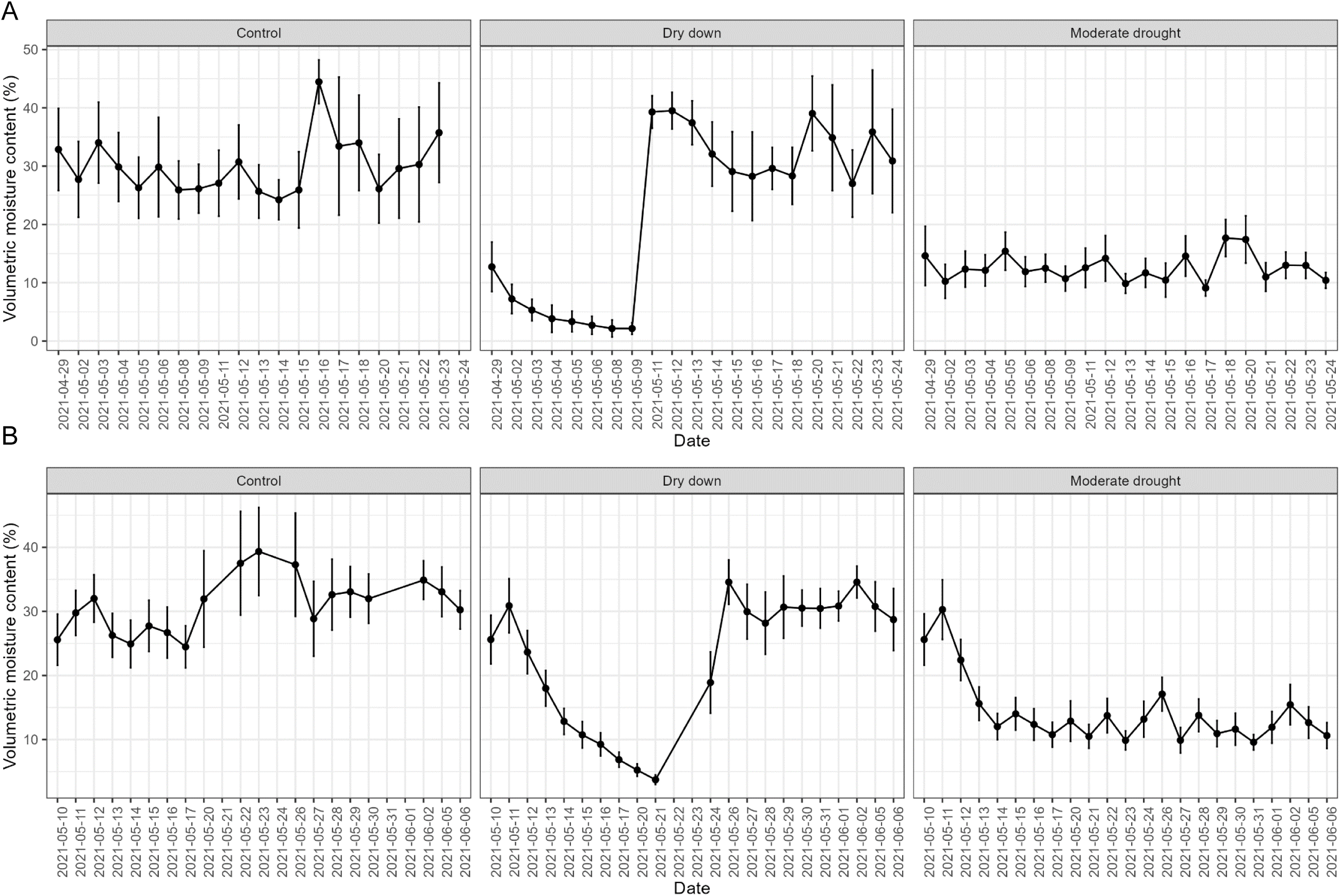
Volumetric moisture content of the pots during the (A) first and (B) second experimental repetition.

**Figure S2:**
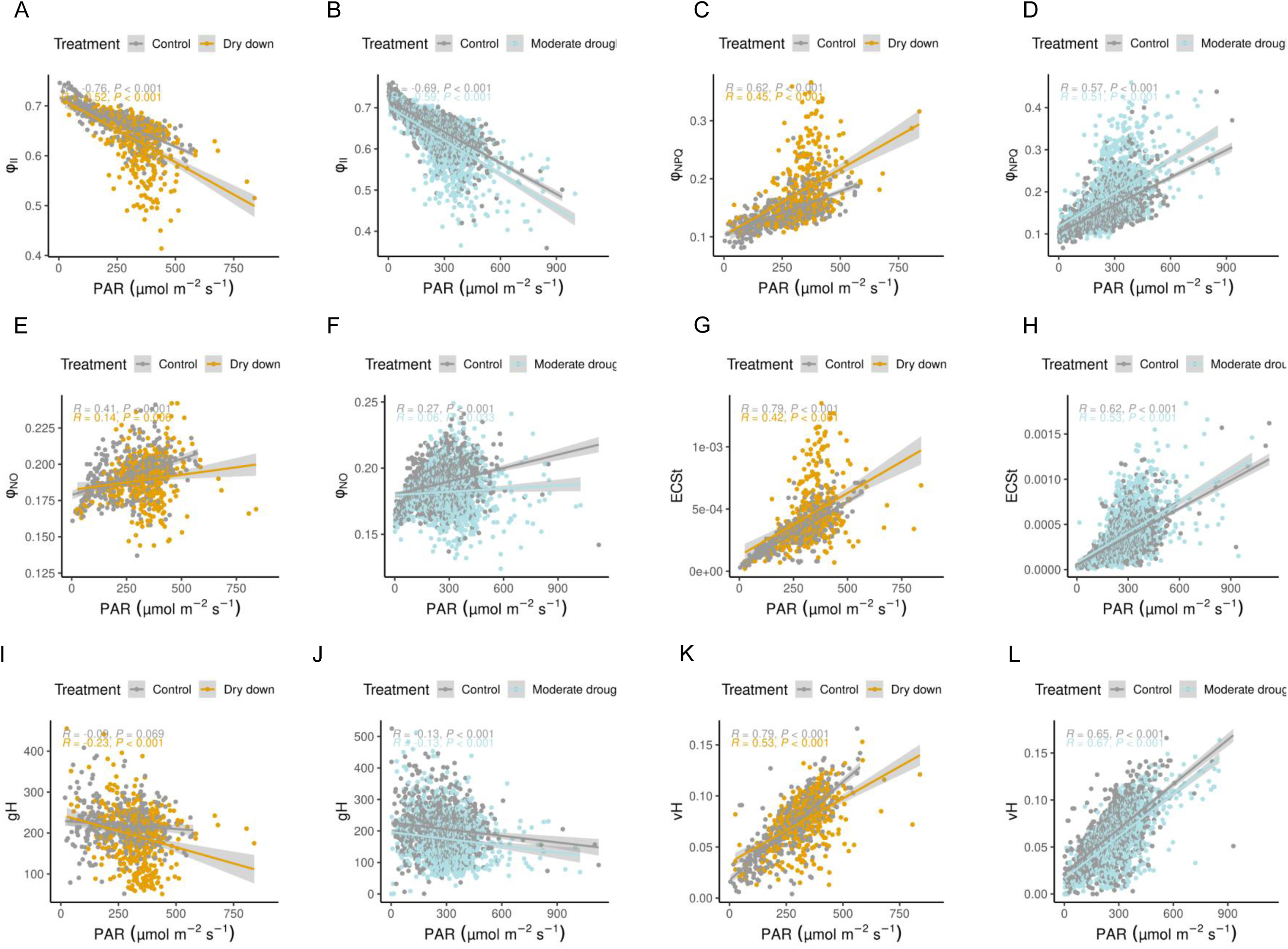
Effect of photosynthetic active radiation (PAR) on photosynthetic parameters compared between control and the two drought stress types. The correlation coefficient between PAR and (A-B) quantum yield of photosystem II (Φ_II_), (C-D) non-photochemical quenching of chlorophyll fluorescence (Φ_NPQ_), (E-F) quantum yield of non-regulatory energy dissipation (Φ_NO_), (G-H) magnitude of electrochromic shift (ECS_t_), (I-J) proton conductivity (gH^+^), and (K-L) steady state proton flux (vH^+^).

**Figure S3:**
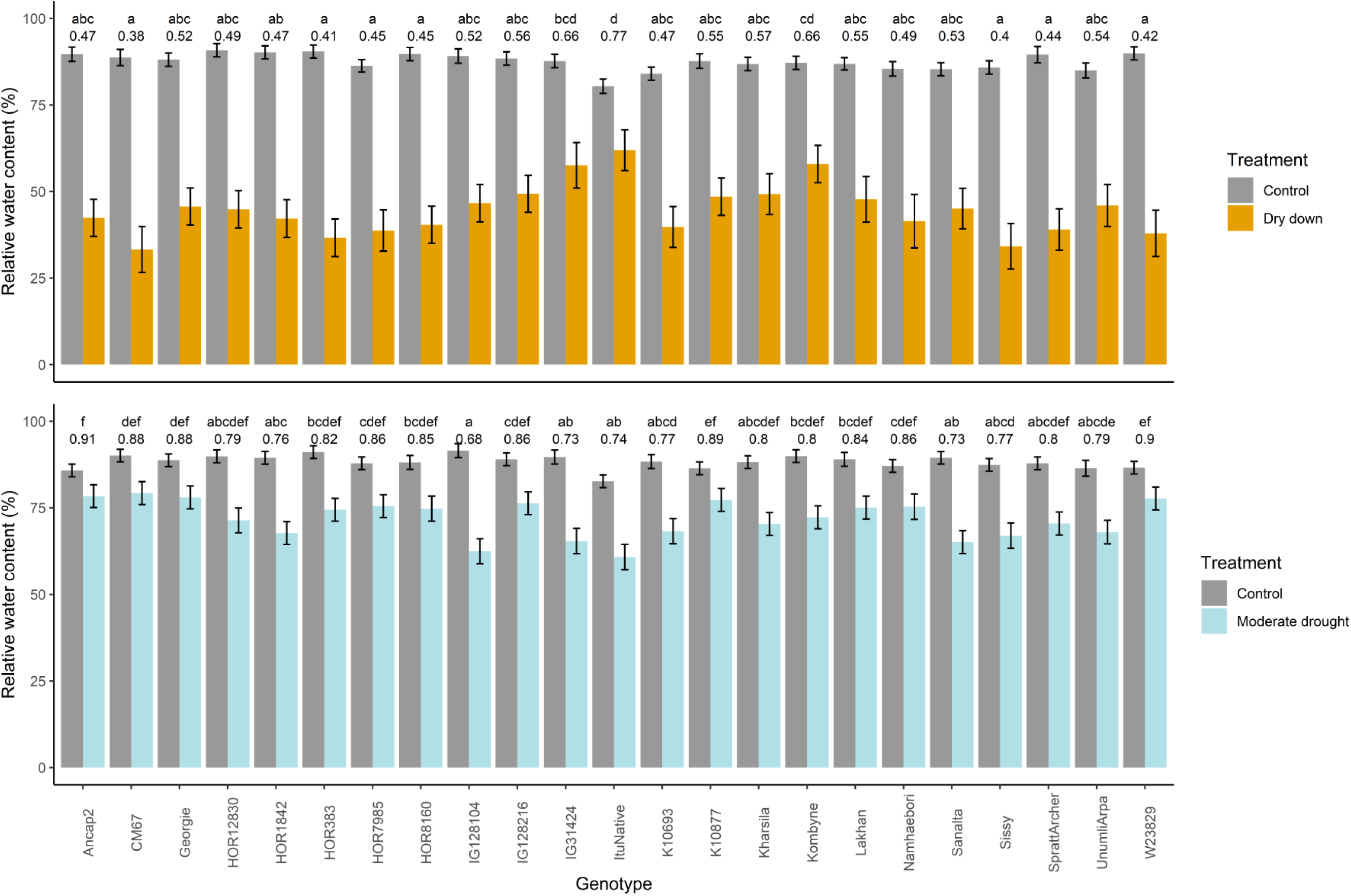
Relative water content (%) under control condition and the two types of drought stress. Drought stress was applied to two-week-old plants. For the dry down treatment, the pots underwent controlled dehydration, and the data were collected on 7 d after the start of drought stress treatment. For moderate drought stress plants were grown under constant volumetric moisture content of 15% and the data were collected on 12 d exposition to moderate drought stress. The experiment was repeated two times with three independent biological replicates per genotype. AEM of the morphological and spectral traits were estimated across two independent experiments. The bar represents adjusted entry mean ± standard error. The number above the bar is the stress response index (SRI). The genotypes not sharing the same letters are significantly different (p < 0.05) for SRI using the Tukey post hoc test.

**Figure S4:**
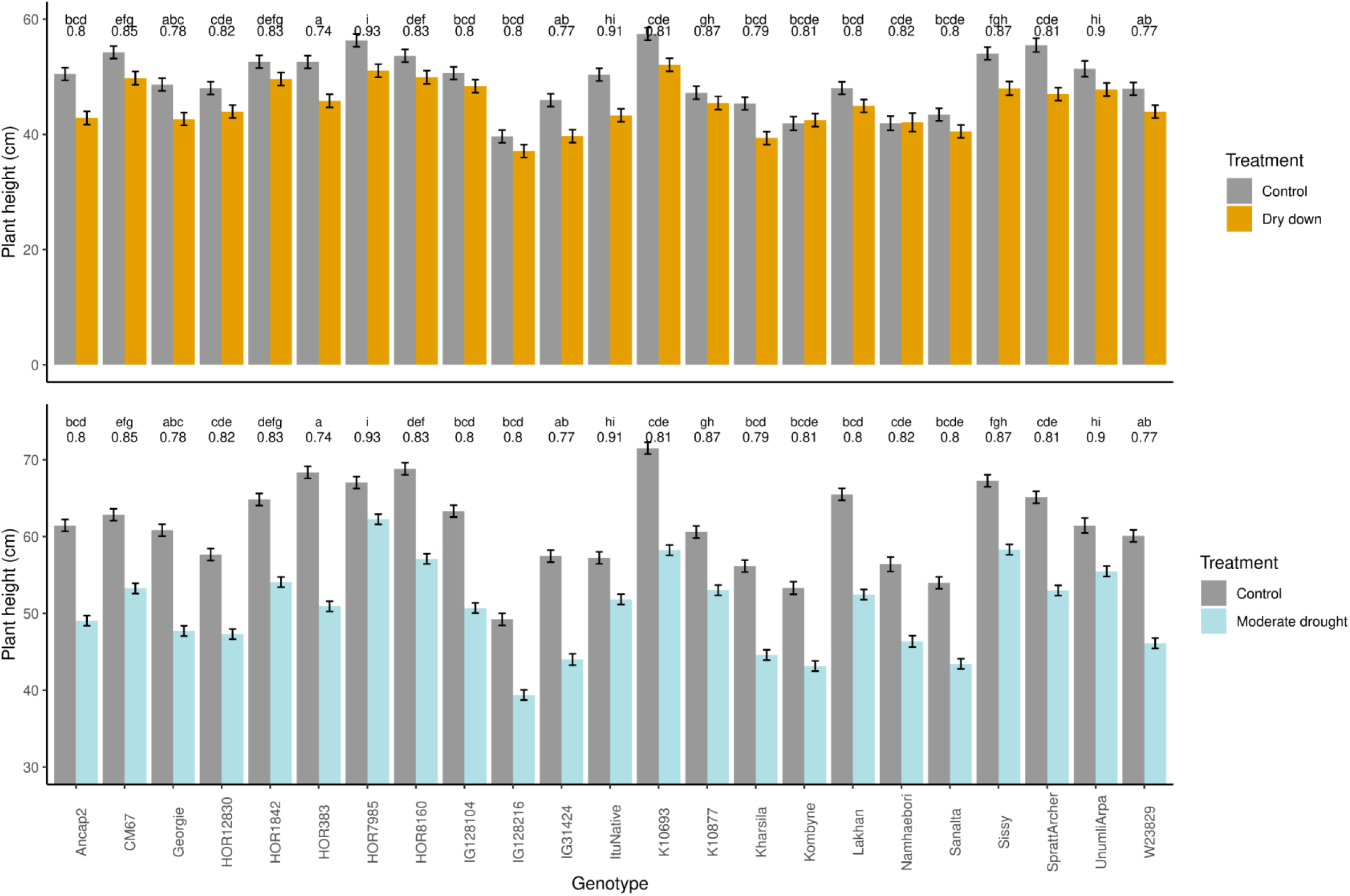
Plant height under control condition and the two types of drought stress. Drought stress was applied to two-week-old plants. For the dry down treatment, the pots underwent controlled dehydration, and the data were collected for 1 d, 4 d and 7 d after the start of drought stress treatment. For moderate drought stress plants were grown under constant volumetric moisture content of 15% and the data were collected from 1 d, 4 d, 7 d, 11 d, 14 d, 18 d, 21 d, 25 d and 28 d after exposition to moderate drought stress. The experiment was repeated two times with three independent biological replicates per genotype. AEM of the morphological and spectral traits were estimated across the time points and two independent experiments. The bar represents adjusted entry mean ± standard error. The number above the bar is the stress response index (SRI). The genotypes not sharing the same letters are significantly different (p < 0.05) for SRI using the Tukey post hoc test.

**Figure S5:**
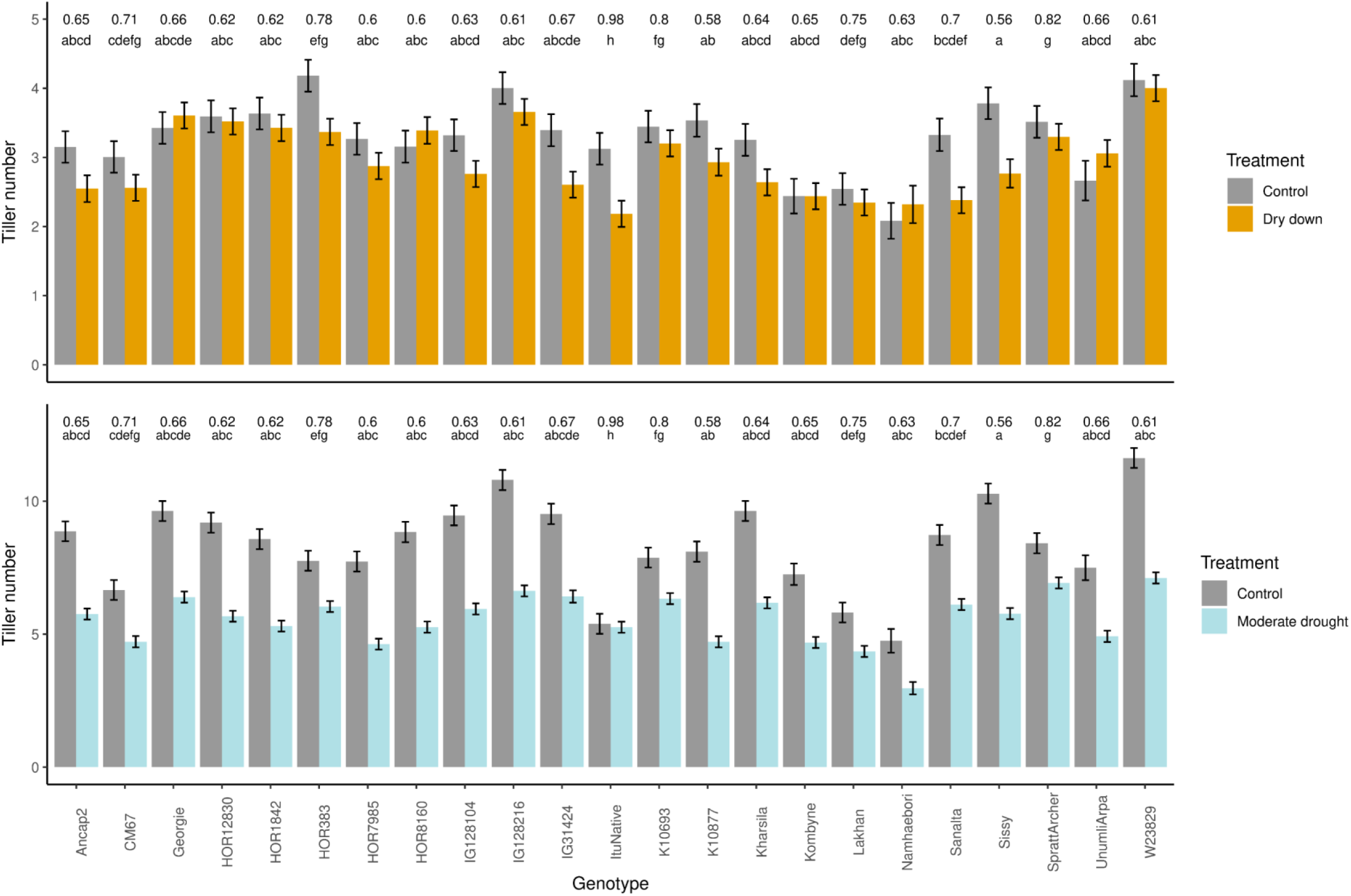
Tiller number under control condition and the two types of drought stress. Drought stress was applied to two-week-old plants. For the dry down treatment, the pots underwent controlled dehydration, and the data were collected for 1 d, 4 d and 7 d after the start of drought stress treatment. For moderate drought stress plants were grown under constant volumetric moisture content of 15% and the data were collected from 1 d, 4 d, 7 d, 11 d, 14 d, 18 d, 21 d, 25 d and 28 d after exposition to moderate drought stress. The experiment was repeated two times with three independent biological replicates per genotype. AEM of the morphological and spectral traits were estimated across the time points and two independent experiments. The bar represents adjusted entry mean ± standard error. The number above the bar is the stress response index (SRI). The genotypes not sharing the same letters are significantly different (p < 0.05) for SRI using the Tukey post hoc test.

**Figure S6:**
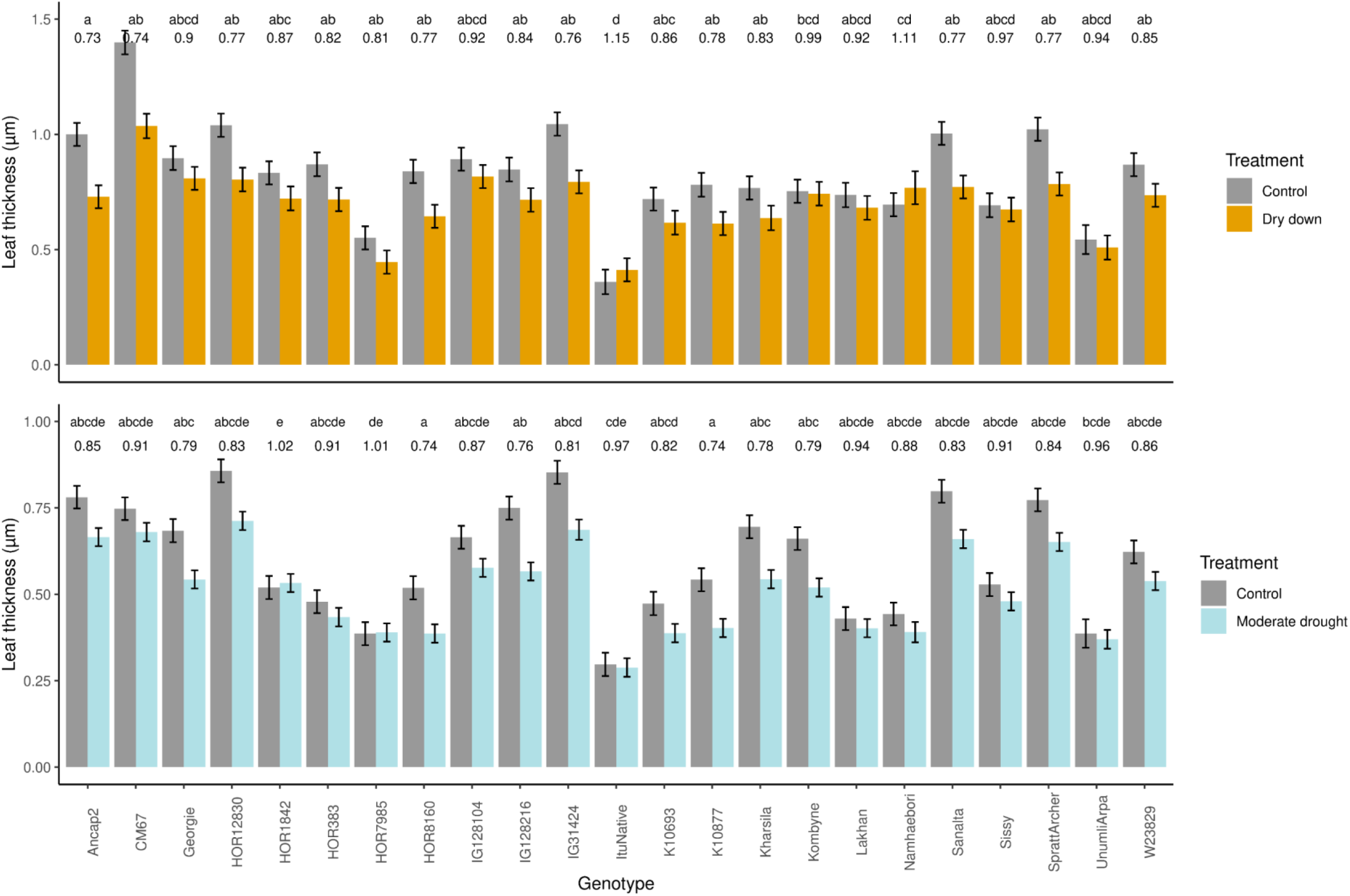
Leaf thickness under control condition and the two types of drought stress. Drought stress was applied to two-week-old plants. For the dry down treatment, the pots underwent controlled dehydration, and the data were collected for 1 d, 4 d and 7 d after the start of drought stress treatment. For moderate drought stress plants were grown under constant volumetric moisture content of 15% and the data were collected from 1 d, 4 d, 7 d, 11 d, 14 d, 18 d, 21 d, 25 d and 28 d after exposition to moderate drought stress. The experiment was repeated two times with three independent biological replicates per genotype. AEM of the morphological and spectral traits were estimated across the time points and two independent experiments. The bar represents adjusted entry mean ± standard error. The number above the bar is the stress response index (SRI). The genotypes not sharing the same letters are significantly different (p < 0.05) for SRI using the Tukey post hoc test.

**Figure S7:**
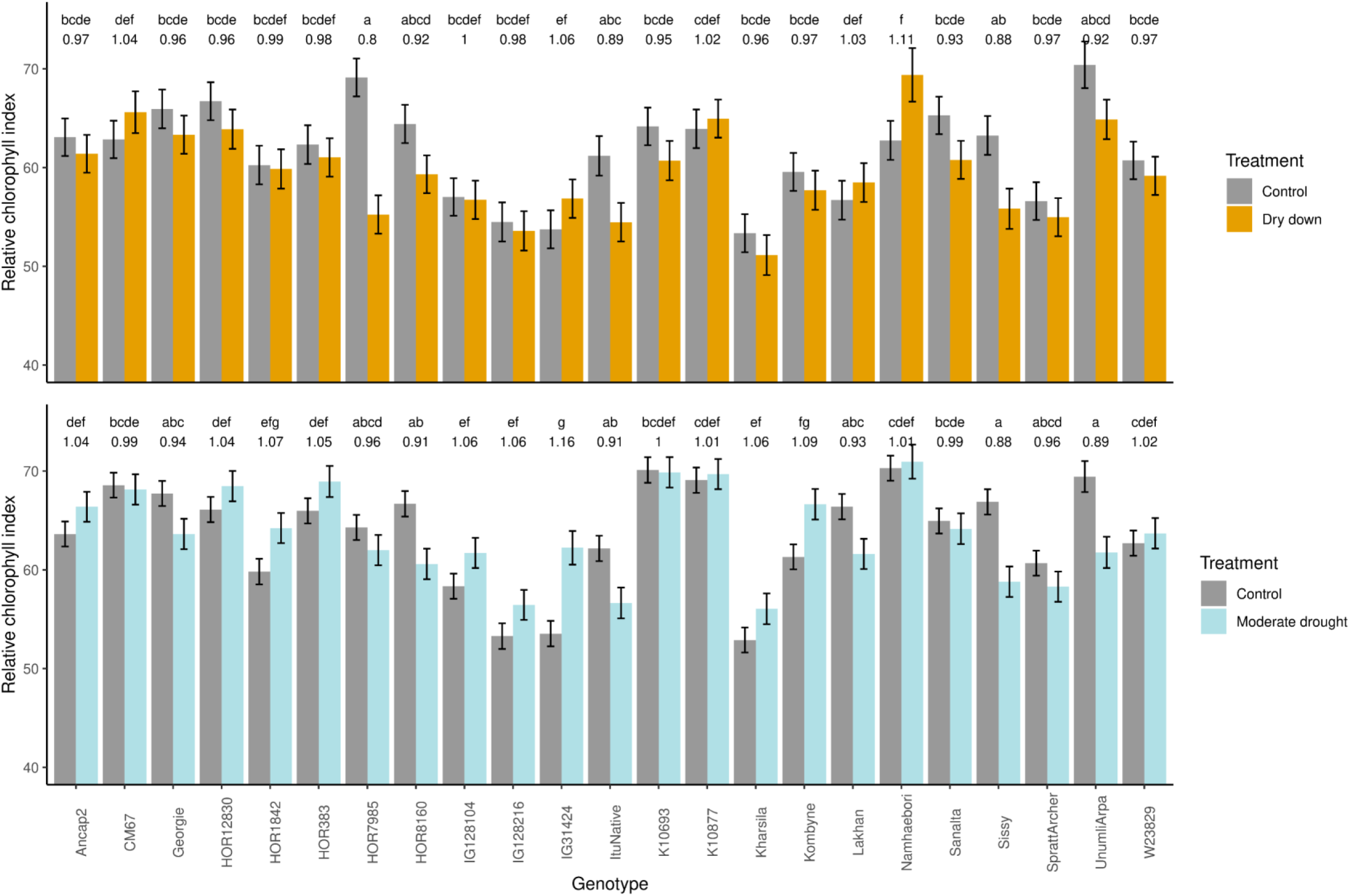
Relative chlorophyll index under control condition and the two types of drought stress. Drought stress was applied to two-week-old plants. For the dry down treatment, the pots underwent controlled dehydration, and the data were collected for 1 d, 4 d and 7 d after the start of drought stress treatment. For moderate drought stress plants were grown under constant volumetric moisture content of 15% and the data were collected from 1 d, 4 d, 7 d, 11 d, 14 d, 18 d, 21 d, 25 d and 28 d after exposition to moderate drought stress. The experiment was repeated two times with three independent biological replicates per genotype. AEM of the morphological and spectral traits were estimated across the time points and two independent experiments. The bar represents adjusted entry mean ± standard error. The number above the bar is the stress response index (SRI). The genotypes not sharing the same letters are significantly different (p < 0.05) for SRI using the Tukey post hoc test.

**Figure S8:**
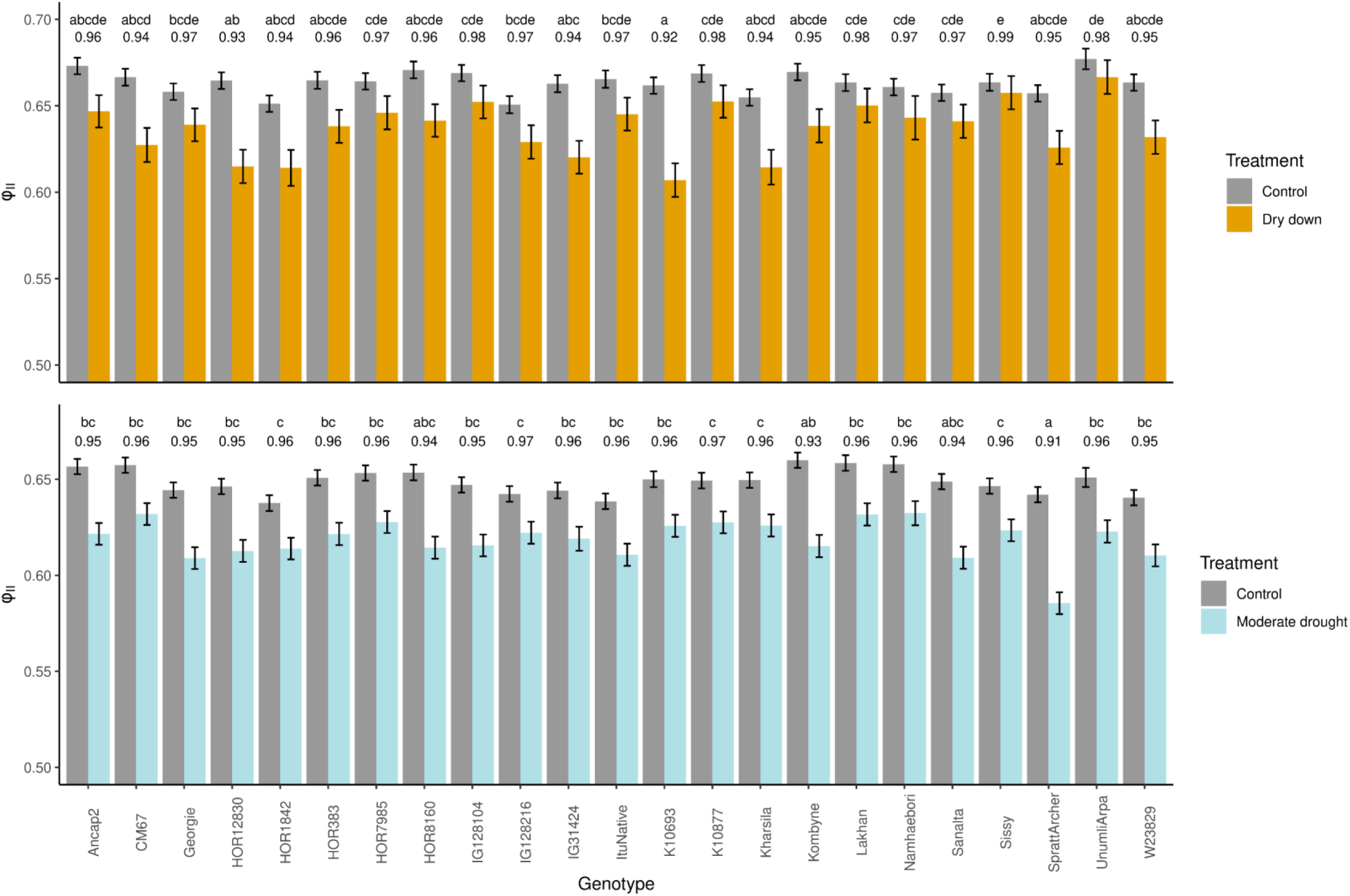
Quantum yield of photosystem II (Φ_II_) under control and water stress conditions. Water stress was applied to two-week-old plants control condition and the two types of drought stress. Drought stress was applied to two-week-old plants. For the dry down treatment, the pots underwent controlled dehydration, and the data were collected for 1 d, 4 d and 7 d after the start of drought stress treatment. For moderate drought stress plants were grown under constant volumetric moisture content of 15% and the data were collected from 1 d, 4 d, 7 d, 11 d, 14 d, 18 d, 21 d, 25 d and 28 d after exposition to moderate drought stress. The experiment was repeated two times with three independent biological replicates per genotype. AEM of the morphological and spectral traits were estimated across the time points and two independent experiments. The bar represents adjusted entry mean ± standard error. The number above the bar is the stress response index (SRI). The genotypes not sharing the same letters are significantly different (p < 0.05) for SRI using the Tukey post hoc test.

**Figure S9:**
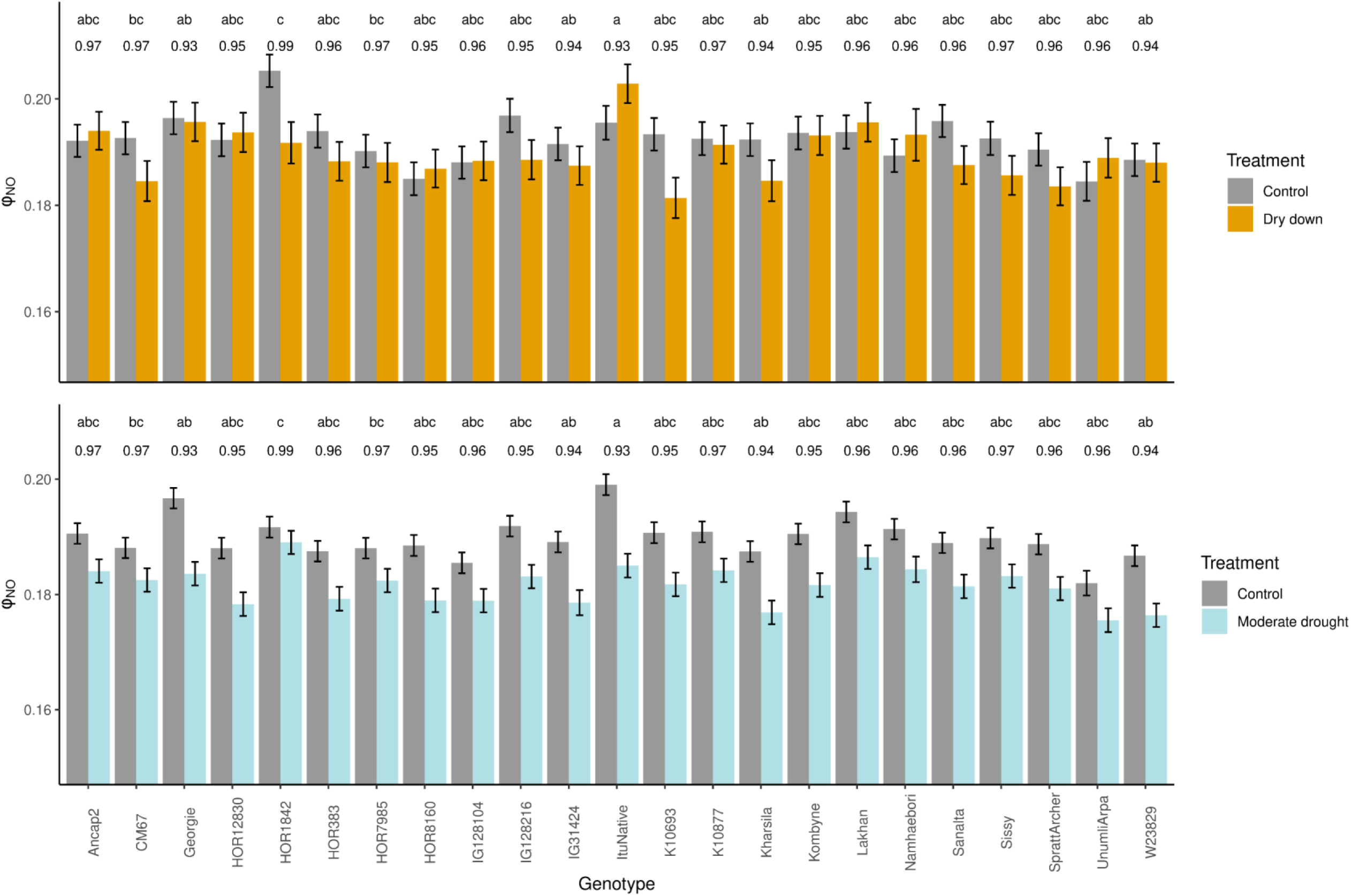
Quantum yield of non-regulatory energy dissipation (Φ_NO_) control condition and the two types of drought stress. Drought stress was applied to two-week-old plants. For the dry down treatment, the pots underwent controlled dehydration, and the data were collected for 1 d, 4 d and 7 d after the start of drought stress treatment. For moderate drought stress plants were grown under constant volumetric moisture content of 15% and the data were collected from 1 d, 4 d, 7 d, 11 d, 14 d, 18 d, 21 d, 25 d and 28 d after exposition to moderate drought stress. The experiment was repeated two times with three independent biological replicates per genotype. AEM of the morphological and spectral traits were estimated across the time points and two independent experiments. The bar represents adjusted entry mean ± standard error. The number above the bar is the stress response index (SRI). The genotypes not sharing the same letters are significantly different (p < 0.05) for SRI using the Tukey post hoc test.

**Figure S10:**
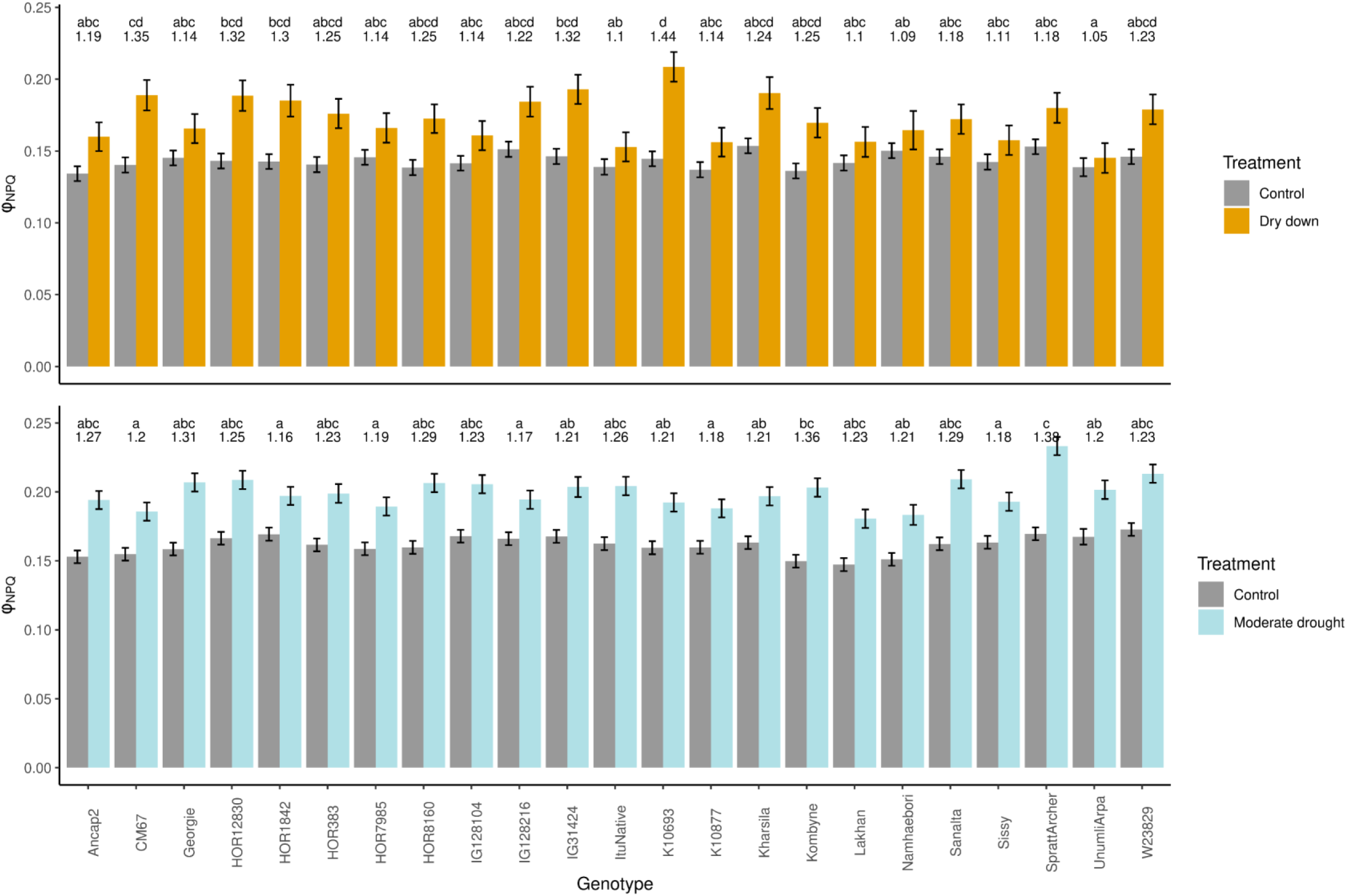
Non-photochemical quenching of chlorophyll fluorescence in photosystem II (Φ_NPQ_) control condition and the two types of drought stress. Drought stress was applied to two-week-old plants. For the dry down treatment, the pots underwent controlled dehydration, and the data were collected for 1 d, 4 d and 7 d after the start of drought stress treatment. For moderate drought stress plants were grown under constant volumetric moisture content of 15% and the data were collected from 1 d, 4 d, 7 d, 11 d, 14 d, 18 d, 21 d, 25 d and 28 d after exposition to moderate drought stress. The experiment was repeated two times with three independent biological replicates per genotype. AEM of the morphological and spectral traits were estimated across the time points and two independent experiments. The bar represents adjusted entry mean ± standard error. The number above the bar is the stress response index (SRI). The genotypes not sharing the same letters are significantly different (p < 0.05) for SRI using the Tukey post hoc test.

**Figure S11:**
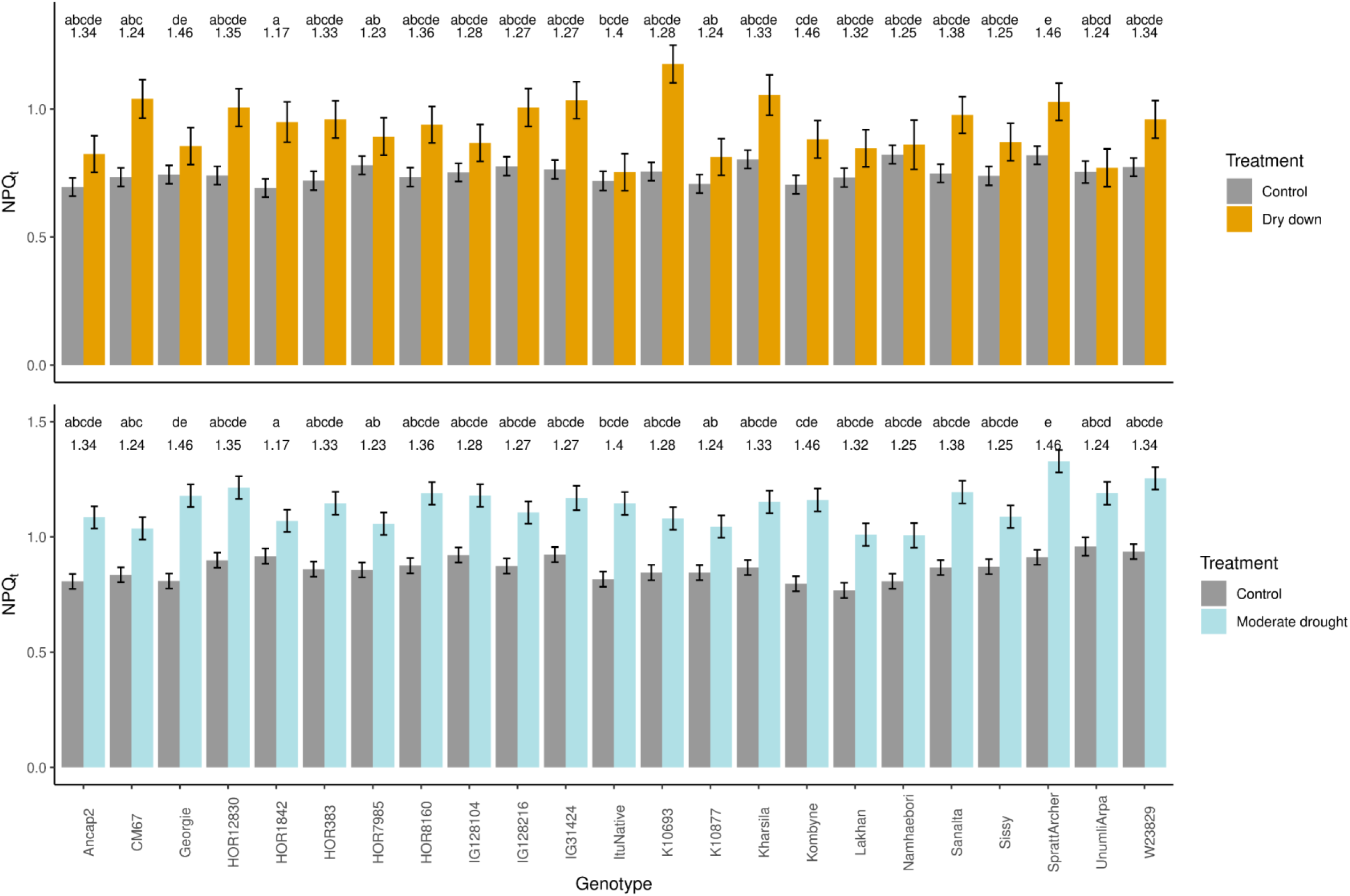
Non-photochemical quenching of chlorophyll fluorescence in photosystem II (NPQ_t_) control condition and the two types of drought stress. Drought stress was applied to two-week-old plants. For the dry down treatment, the pots underwent controlled dehydration, and the data were collected for 1 d, 4 d and 7 d after the start of drought stress treatment. For moderate drought stress plants were grown under constant volumetric moisture content of 15% and the data were collected from 1 d, 4 d, 7 d, 11 d, 14 d, 18 d, 21 d, 25 d and 28 d after exposition to moderate drought stress. The experiment was repeated two times with three independent biological replicates per genotype. AEM of the morphological and spectral traits were estimated across the time points and two independent experiments. The bar represents adjusted entry mean ± standard error. The number above the bar is the stress response index (SRI). The genotypes not sharing the same letters are significantly different (p < 0.05) for SRI using the Tukey post hoc test.

**Figure S12:**
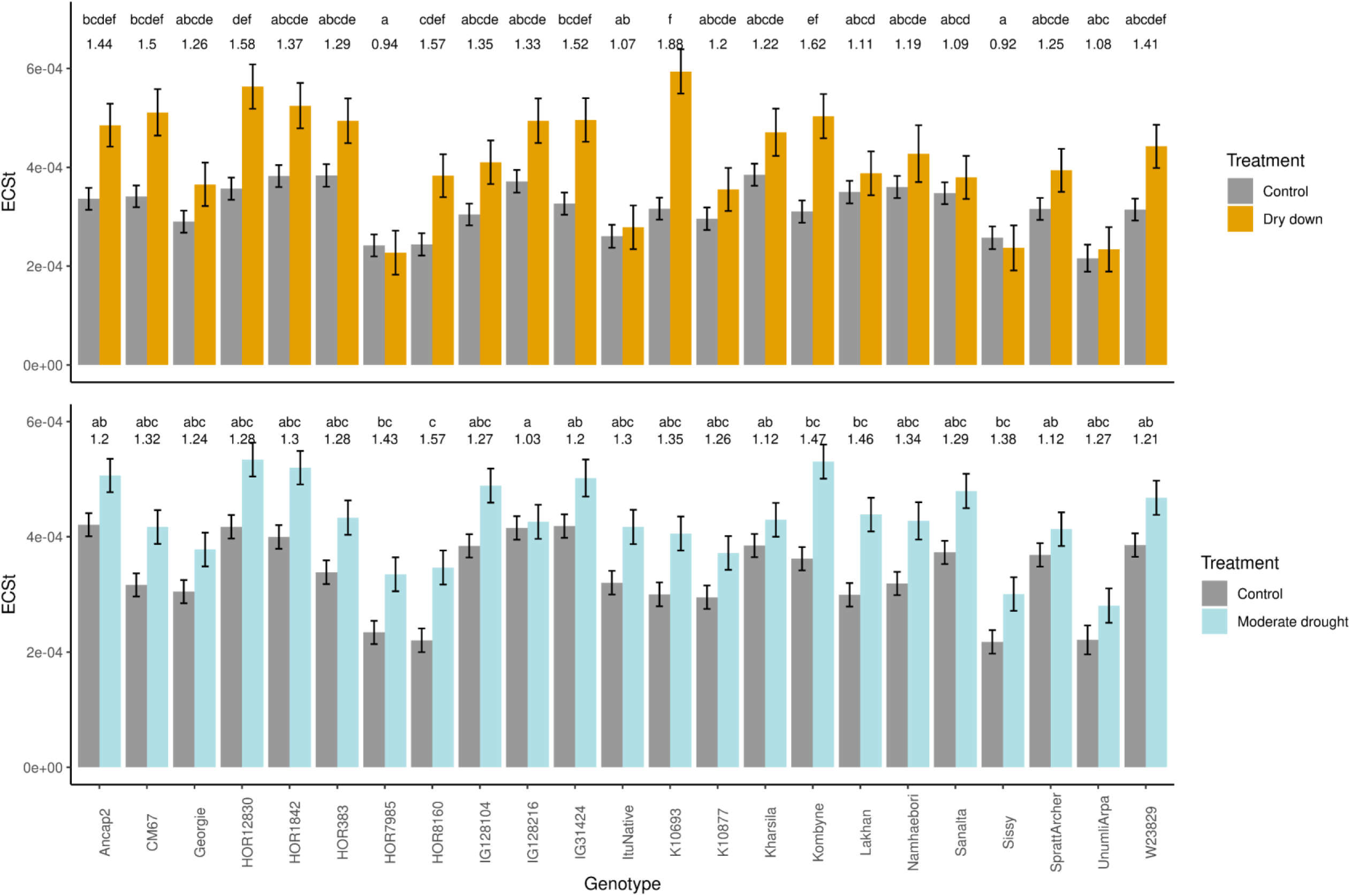
Magnitude of electrochromic shift (ECS_t_) under control condition and the two types of drought stress. Drought stress was applied to two-week-old plants. For the dry down treatment, the pots underwent controlled dehydration, and the data were collected for 1 d, 4 d and 7 d after the start of drought stress treatment. For moderate drought stress plants were grown under constant volumetric moisture content of 15% and the data were collected from 1 d, 4 d, 7 d, 11 d, 14 d, 18 d, 21 d, 25 d and 28 d after exposition to moderate drought stress. The experiment was repeated two times with three independent biological replicates per genotype. AEM of the morphological and spectral traits were estimated across the time points and two independent experiments. The bar represents adjusted entry mean ± standard error. The number above the bar is the stress response index (SRI). The genotypes not sharing the same letters are significantly different (p < 0.05) for SRI using the Tukey post hoc test.

**Figure S13:**
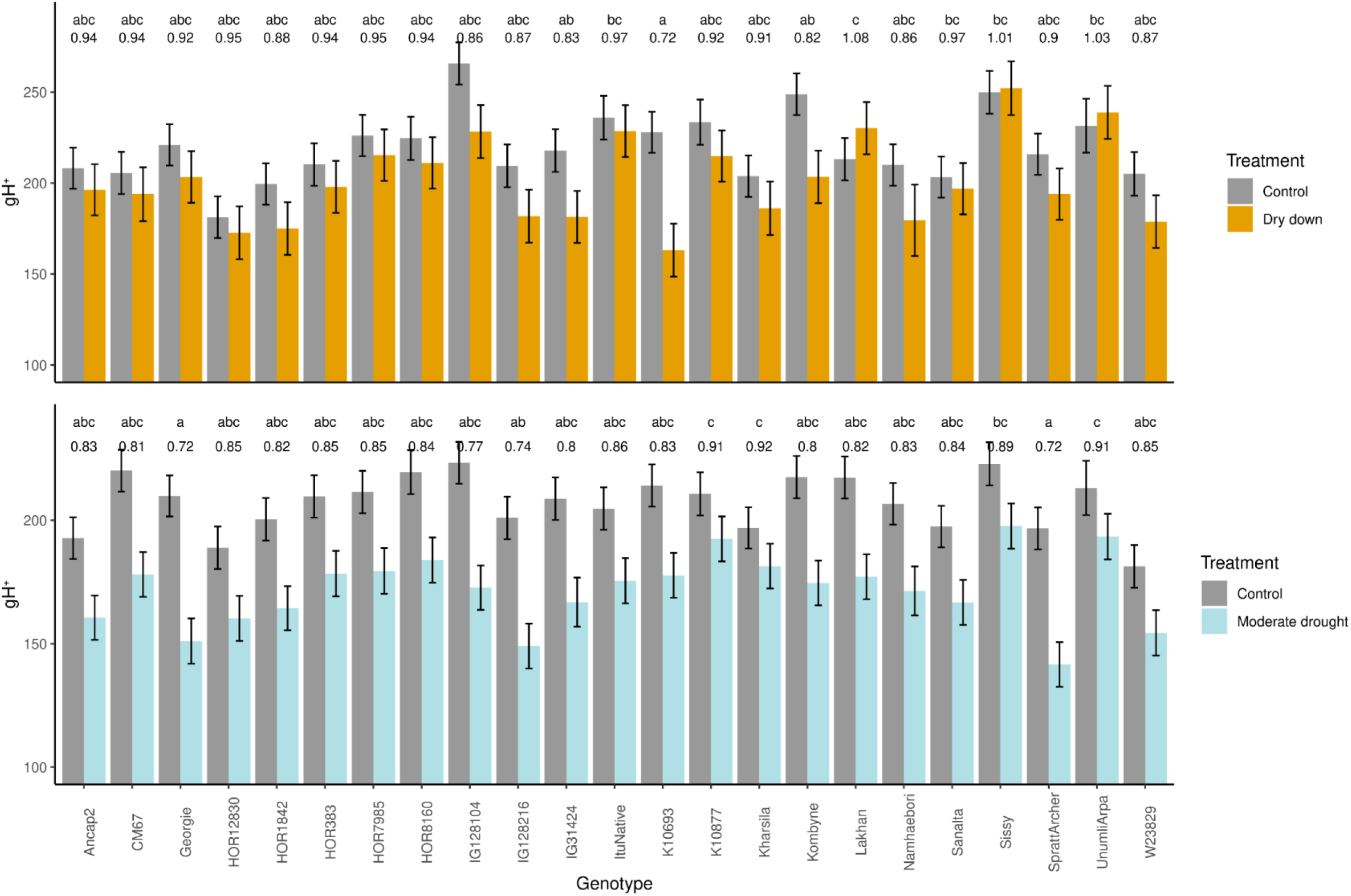
Proton conductivity (gH^+^) under control condition and the two types of drought stress. Drought stress was applied to two-week-old plants. For the dry down treatment, the pots underwent controlled dehydration, and the data were collected for 1 d, 4 d and 7 d after the start of drought stress treatment. For moderate drought stress plants were grown under constant volumetric moisture content of 15% and the data were collected from 1 d, 4 d, 7 d, 11 d, 14 d, 18 d, 21 d, 25 d and 28 d after exposition to moderate drought stress. The experiment was repeated two times with three independent biological replicates per genotype. AEM of the morphological and spectral traits were estimated across the time points and two independent experiments. The bar represents adjusted entry mean ± standard error. The number above the bar is the stress response index (SRI). The genotypes not sharing the same letters are significantly different (p < 0.05) for SRI using the Tukey post hoc test.

**Figure S14:**
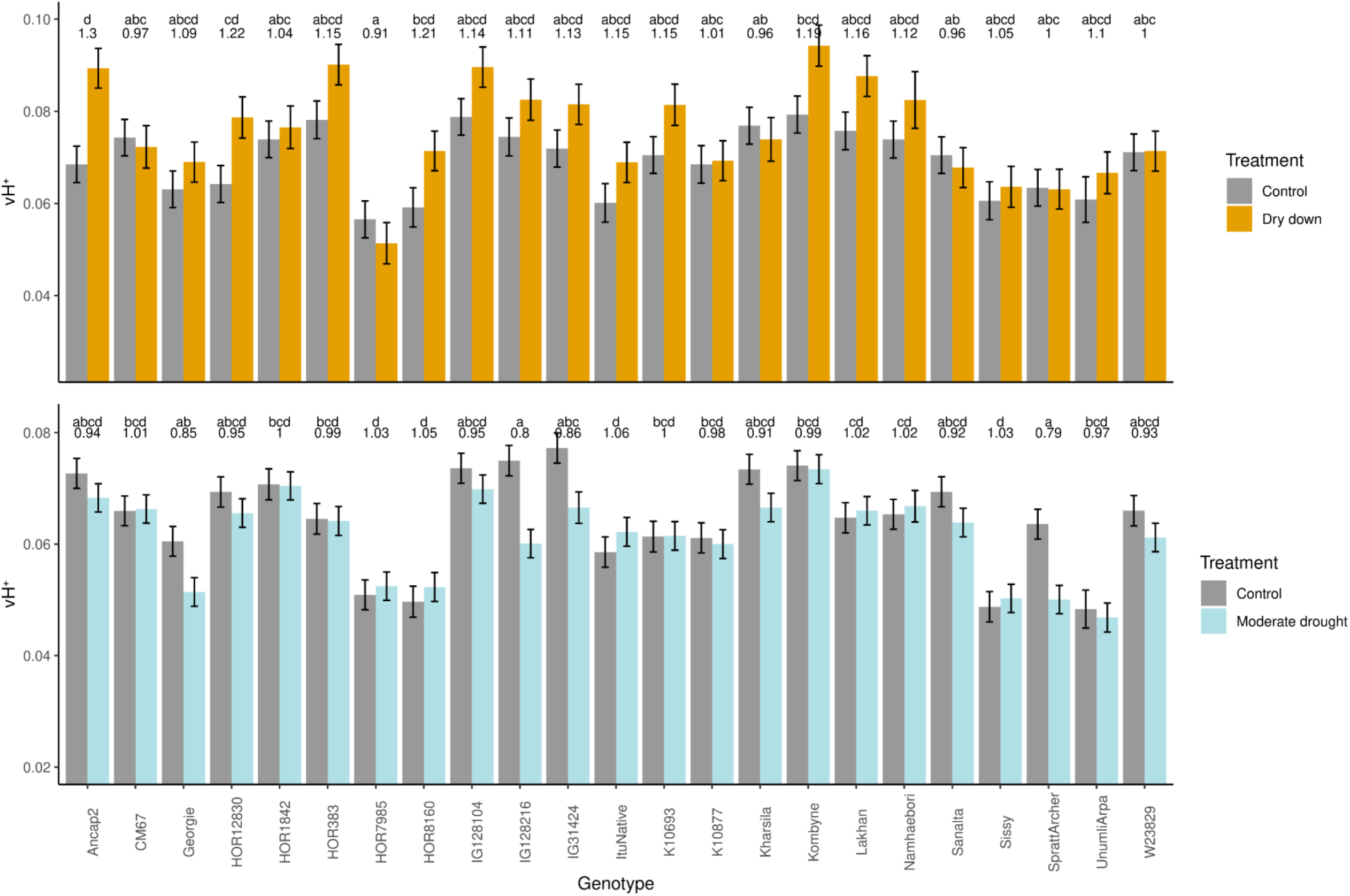
Steady state proton flux (vH^+^) under control condition and the two types of drought stress. Drought stress was applied to two-week-old plants. For the dry down treatment, the pots underwent controlled dehydration, and the data were collected for 1 d, 4 d and 7 d after the start of drought stress treatment. For moderate drought stress plants were grown under constant volumetric moisture content of 15% and the data were collected from 1 d, 4 d, 7 d, 11 d, 14 d, 18 d, 21 d, 25 d and 28 d after exposition to moderate drought stress. The experiment was repeated two times with three independent biological replicates per genotype. AEM of the morphological and spectral traits were estimated across the time points and two independent experiments. The bar represents adjusted entry mean ± standard error. The number above the bar is the stress response index (SRI). The genotypes not sharing the same letters are significantly different (p < 0.05) for SRI using the Tukey post hoc test.

**Figure S15:**
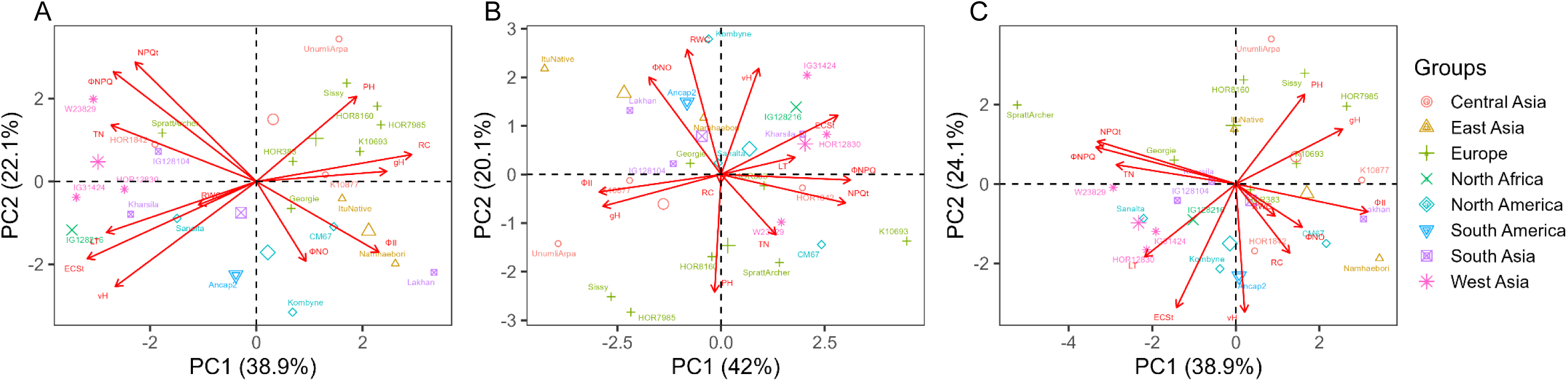
Principal component (PC) analysis of morphological, physiological and photosynthetic parameters. Adjusted entry mean (AEM) of evaluated parameters was used to perform principal component analysis for (A) control conditions (B) dry down stress (C) moderate drought stress. The points represent genotypes and the vectors represent the traits namely tiller numbers (TN), plant height (PH), leaf thickness (LT), relative chlorophyll (RC), relative water content (RWC), quantum yield of photosystem II (Φ_II_), non-photochemical losses (Φ_NO_), the fraction of absorbed light dissipated by non-photochemical quenching(Φ_NPQ_ and NPQ_t_), magnitude of electrochromic shift (ECS_t_), proton conductivity (gH^+^) and steady state proton flux (vH^+^). For the dry down treatment, the pots underwent controlled dehydration, and the data were collected for 1 d, 4 d and 7 d after the start of drought stress treatment. For moderate drought stress plants were grown under constant volumetric moisture content of 15% and the data were collected from 1 d, 4 d, 7 d, 11 d, 14 d, 18 d, 21 d, 25 d and 28 d after exposition to MD stress. The experiment was repeated two times with three independent biological replicates per genotype. AEM of the evaluated parameters was estimated across the time points and two independent experiments.

**Figure S16:**
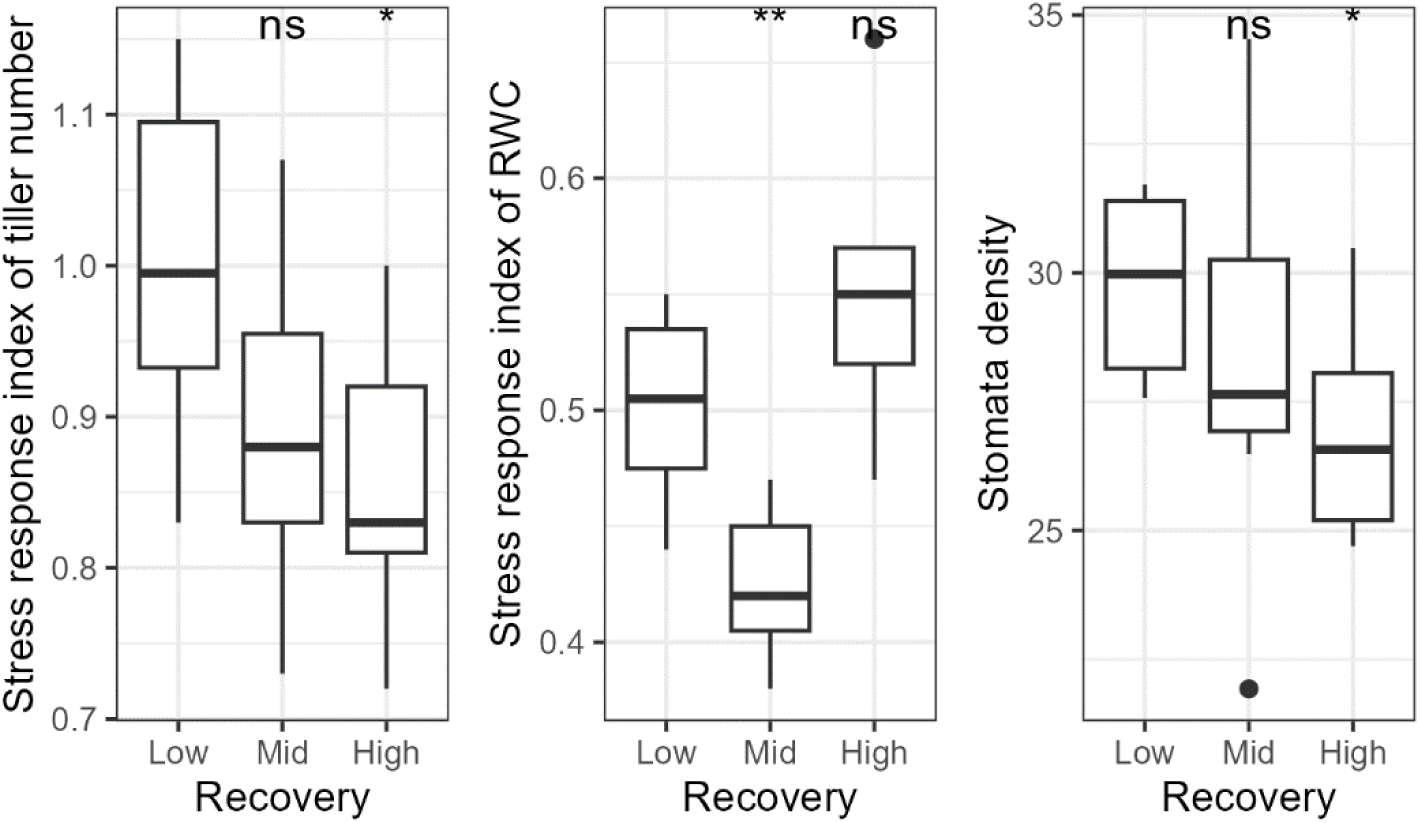
Association between count of recovered plants and morphological and physiological parameters. The distribution of stress response index calculated as the ratio of adjusted entry mean (AEM) for (A) tiller number (TN) and (B) relative water content observed in dry down stress versus control conditions; and (C) AEM of stomata density among inbreds that showed low, medium and high recovery count after stress releases. For the dry down stress, the pots underwent controlled dehydration, and the data were collected for 1 d, 4 d and 7 d after the start of drought stress treatment. The experiment was repeated two times with three independent biological replicates per genotype. AEM was estimated across the time points and two independent experiments. Plants were rewatered 12 d after the start of dry down stress. Then, the plants that produced true leaves and resumed growth two weeks after the start of rewatering were counted as recovered plants. AEM of stomata density were estimated across two independent experiments at three different leaf position from two abaxial and adaxial surface of leaf blade. Asterisks indicate significant difference between reference group (inbreds with low recovery count) versus other group of inbreds using student’s t-test.

**Figure S17:**
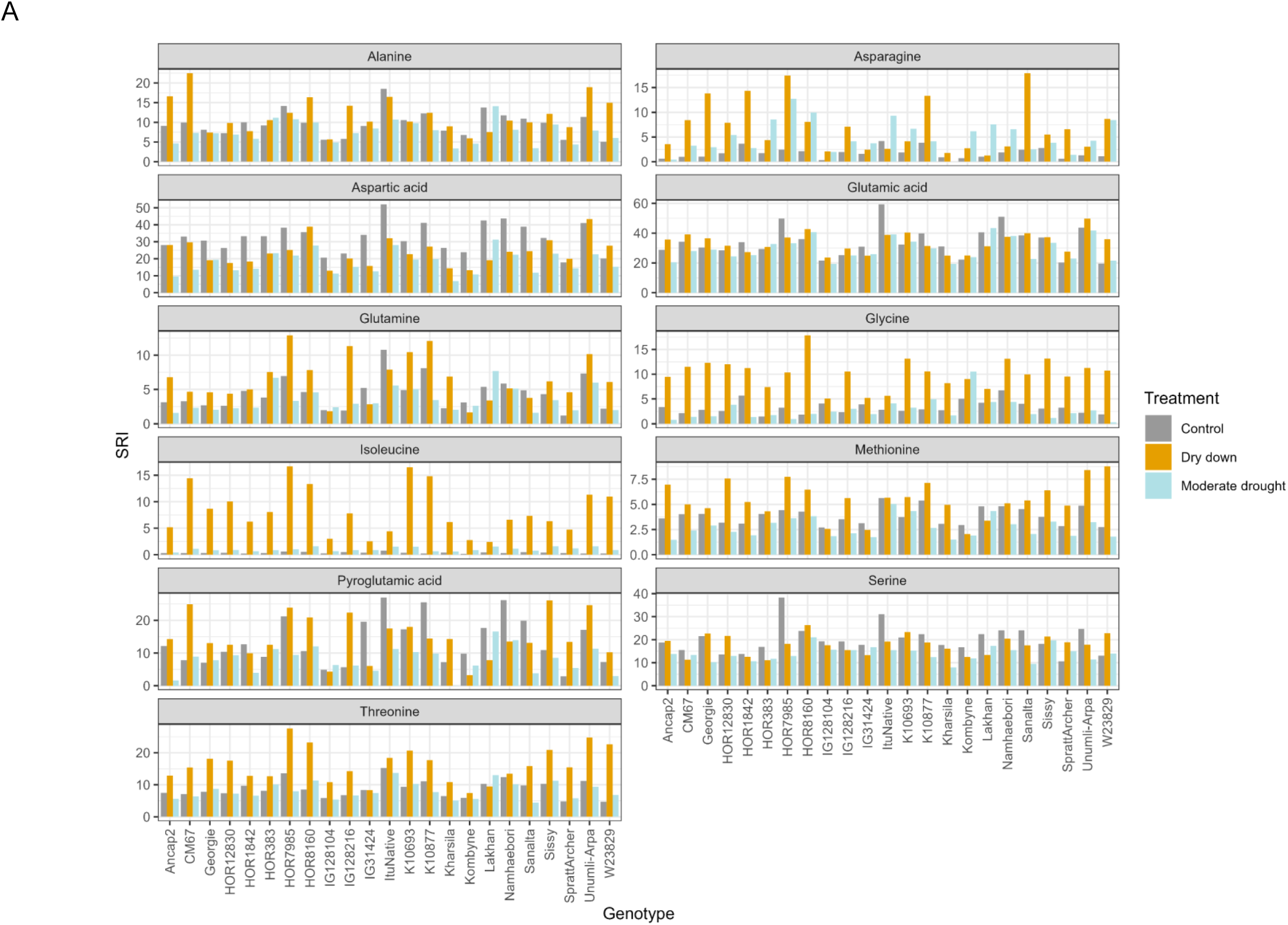
Adjusted entry mean of amino acids under different growing conditions. The metabolite profiling was done in the leaf samples collected from 7 d and 12 d after stress under dry down and moderate drought stress, respectively. The leaves from the control conditions collected at 7 d and 12 d were pooled together making one representative control group. The experiment was repeated two times with three independent biological replicates per genotype. Adjusted entry mean of the metabolites were estimated across two independent experiments.

**Figure S18:**
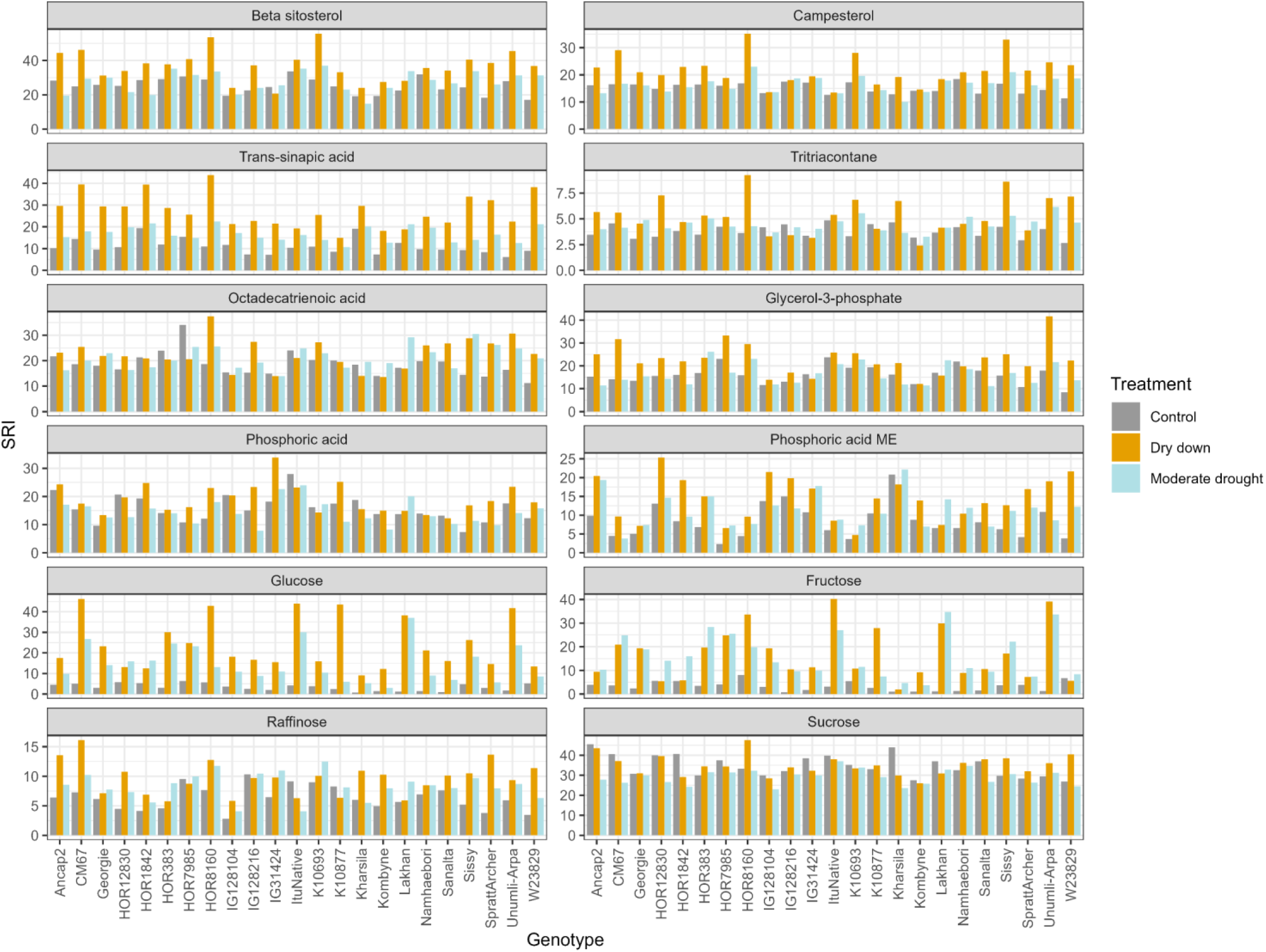
Adjusted entry mean of lipids, sugars, phosphates and phenylpropanoids under different growing conditions. The metabolite profiling was done in the leaf samples collected from 7 d and 12 d after stress under dry down and moderate drought stress, respectively. The leaves from the control conditions collected at 7 d and 12 d were pooled together making one representative control group. The experiment was repeated two times with three independent biological replicates per genotype. Adjusted entry mean of the metabolites were estimated across two independent experiments.

**Figure S19:**
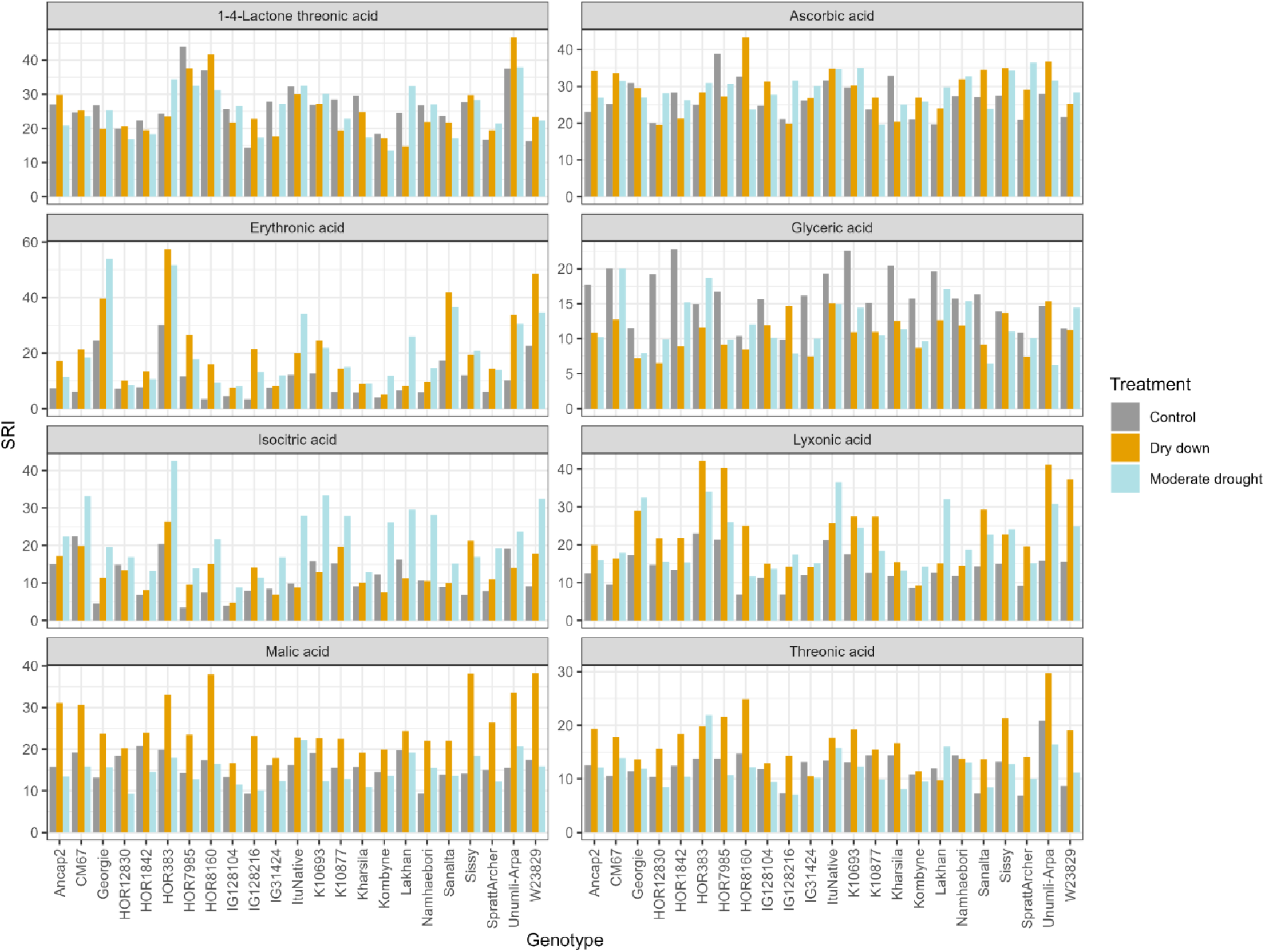
Adjusted entry mean of organic acids under different growing conditions. The metabolite profiling was done in the leaf samples collected from 7 d and 12 d after stress under dry down and moderate drought stress, respectively. The leaves from the control conditions collected at 7 d and 12 d were pooled together making one representative control group. The experiment was repeated two times with three independent biological replicates per genotype. Adjusted entry mean of the metabolites were estimated across two independent experiments.

**Figure S20:**
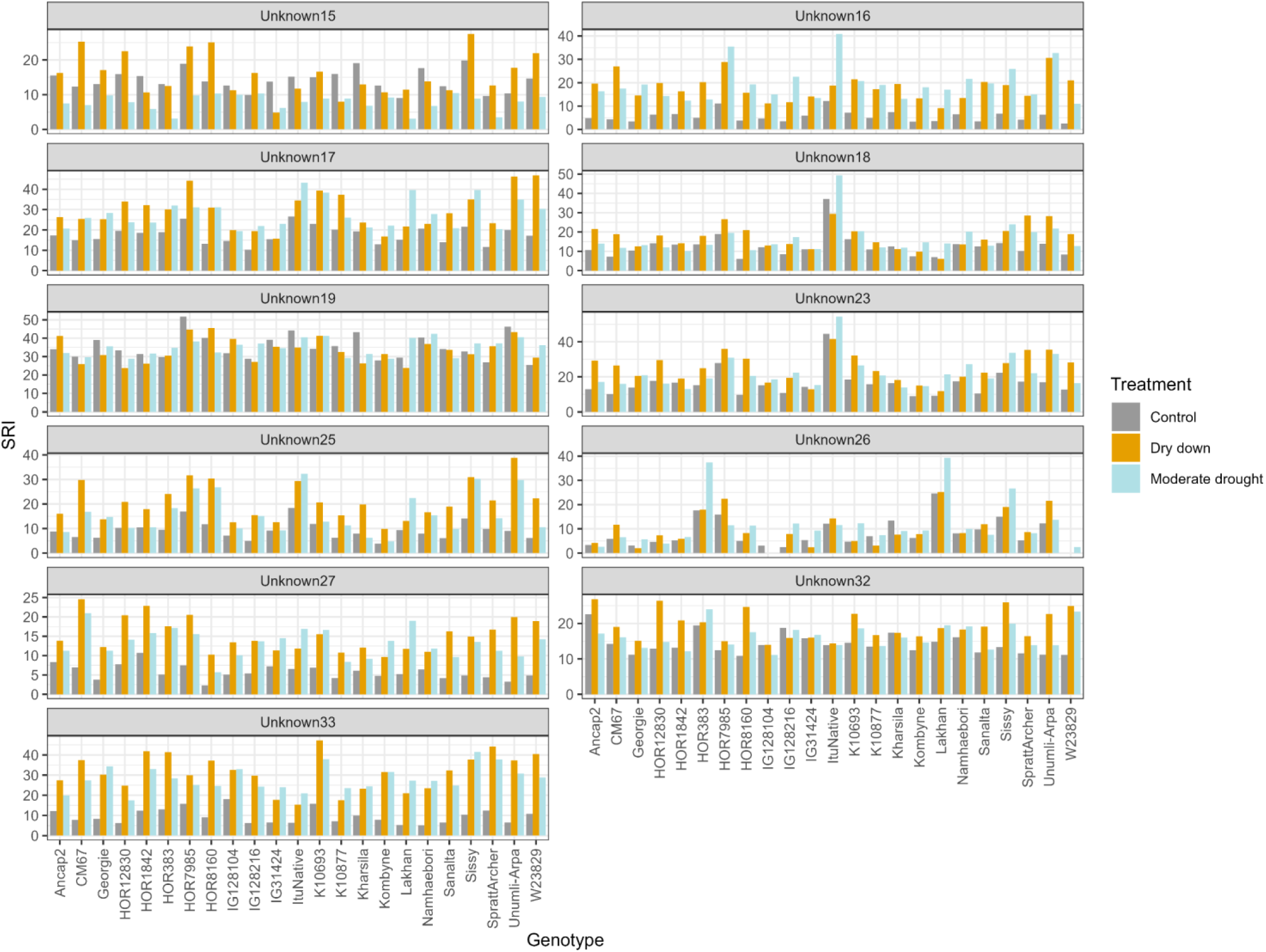
Adjusted entry mean of unknown metabolites under different growing conditions. The metabolite profiling was done in the leaf samples collected from 7 d and 12 d after stress under dry down and moderate drought stress, respectively. The leaves from the control conditions collected at 7 d and 12 d were pooled together making one representative control group. The experiment was repeated two times with three independent biological replicates per genotype. Adjusted entry mean of the metabolites were estimated across two independent experiments.

